# Activity-resolved microbial community profiling using *rpoB* gene and transcript sequencing

**DOI:** 10.64898/2026.08.25.746930

**Authors:** Fabien Cholet, William T. Sloan, Cindy J. Smith

## Abstract

Determining which members of a microbial community are metabolically active remains a central challenge in microbial ecology. Although the *16S rRNA* gene is the dominant marker for bacterial community profiling, it cannot reliably distinguish active cells from dormant or dead populations. As a result, complementary phylogenetic markers whose transcript abundance more closely reflects cellular activity are needed. Here, we systematically evaluated 80 Bacterial protein-coding marker genes and identified *rpoB*, encoding the β-subunit of bacterial RNA polymerase, as the optimal candidate. We designed a new primer pair (1528F/2041R) from a curated database of 305,274 unique *rpoB* sequences and validated it for quantitative PCR and amplicon sequencing of DNA and RNA templates. The *rpoB* qPCR assay achieved a limit of quantification two orders of magnitude lower than the benchmark *16S rRNA* assay, for which a limit of detection could not be determined because of no-template-control amplification. In soil and sediment communities, *rpoB* recovered community composition comparable to *16S rRNA* while providing a quantitative activity signal: *rpoB* cDNA:DNA ratios correlated significantly with taxon-level transcript abundance (R² = 0.22–0.29, p < 0.001), whereas *16S rRNA* ratios did not (p > 0.5). In a biological activated carbon biofilter experiment, *rpoB* transcript abundance tracked the decline in dissolved organic carbon removal rates across a 72-hour time series (correlation coefficients ∼0.84 and 0.99), whereas *16S rRNA* transcripts were uninformative (correlation coefficients between -0.4 and 0.98). These results establish *rpoB* as a quantitatively robust, activity-responsive complement to *16S rRNA* for linking community composition to ecosystem processes.

## 1. Introduction

Microbes and their metabolic activities drive global biogeochemical cycles across natural and engineered environments. Understanding which microorganisms are present and, critically, which are metabolically active is therefore central to linking microbiome composition with ecosystem function. Yet determining which organisms are metabolically active and driving observed process rates remains a fundamental challenge in environmental microbiology. The distinction between presence and activity is not trivial as in natural environments, a large fraction of the microbial community exists in dormant or inactive states, contributing little to ongoing biogeochemical processes despite being numerically abundant (Jones & Lennon, 2010; Lennon & Jones, 2011; Blagodatskaya & Kuzyakov, 2013). Despite its widespread utility, DNA-based approaches provide limited information regarding microbial physiological state. Environmental DNA can persist after cell death, while dormant organisms may remain detectable despite contributing little to ongoing ecosystem processes. To address this limitation, RNA-based approaches have increasingly been used to infer microbial activity, based on the assumption that RNA molecules are generally less stable and more closely linked to ongoing cellular function than DNA (Morgado-Gamero et al., 2025, Rauhut & Klug, 1999).

Quantification and sequencing of reverse-transcribed RNA (cDNA) derived from *16S rRNA* has been widely used in environmental microbiology and have provided important insights into active microbial fractions in soils, sediments, anaerobic digesters, and drinking water systems (Bai et al., 2023, De Vrieze et al., 2018; Hoshino & Matsumoto, 2007; Lillis et al., 2009; Schostag et al., 2015; von Hoyningen-Huene et al., 2022; Yang et al., 2022). However, the use of ribosomal RNA as a proxy for microbial activity remains problematic. Ribosomal RNA molecules are relatively stable compared to messenger RNA transcripts (Li et al., 2017; Morgado-Gamero et al., 2025) and can persist under nutrient limitation or after growth arrest. Furthermore, ribosome abundance is not always tightly coupled to metabolic activity or growth rate, leading to uncertainty when interpreting *16S rRNA* transcript abundance as a measure of microbial activity (Blazewicz et al., 2013; Emerson et al., 2017; Wang et al., 2023). These limitations have prompted growing interest in alternative marker genes that may better reflect transcriptionally active microbial populations.

Single-copy protein-coding housekeeping genes involved in essential cellular functions represent promising complementary markers for microbial community analysis. In contrast to the multicopy *16S rRNA* gene, many protein-coding genes occur as single copies within bacterial genomes, reducing biases associated with variable gene copy number and intragenomic heterogeneity. Moreover, because these genes encode functional cellular processes, their transcripts may more directly reflect ongoing metabolism and growth-related activity (Staeubli et al., 2025). Several conserved protein-coding genes have therefore been proposed as alternative phylogenetic and ecological markers, including *gyrB, recA, rplB,* and *rpoB* (Santos & Ochman 2004, Yamamoto & Harayama 1995, Barret et al., 2015). Among these candidates, the *rpoB* gene, encoding the β-subunit of bacterial RNA polymerase, is particularly attractive because of its central role in transcription. Previous studies have shown that *rpoB*-based approaches can provide improved taxonomic resolution compared with *16S rRNA* (Vos et al., 2012) and recover community structures comparable to shotgun metagenomics (Durand et al., 2025). Evidence has also emerged suggesting that *rpoB* transcripts may better reflect transcriptionally active microbial populations than ribosomal RNA-based approaches (Barnard et al., 2013, 2015). However, despite increasing interest in protein-coding marker genes, several important limitations remain. Existing *rpoB* primers vary considerably in taxonomic coverage, degeneracy, and suitability for quantitative applications. Furthermore, previous studies have typically focused either on taxonomic profiling or on specific environments, and there has been no systematic cross-gene evaluation integrating in silico coverage, quantitative PCR performance, and experimental validation of both DNA and RNA-based community profiling.

To address this, we conducted a comprehensive evaluation of bacterial protein-coding marker genes to identify robust complements to *16S rRNA* for quantitative microbial community profiling and transcriptional activity assessment. We first constructed a curated *rpoB* reference database containing more than 300,000 unique sequences and selected a subset of >20,000 unique sequences from unique species to design and evaluate broad coverage *rpoB* primers suitable for both quantitative PCR and amplicon sequencing. Another seventy-nine candidate marker genes were tested for in-silico primer design and the performance of de-novo and existing primers targeting these other protein markers was compared to the best performing *rpoB* primer pairs. PCR tests were then used to select a single best performing primer pair, targeting the *rpoB* gene. This *rpoB* primer pair was subsequently benchmarked against commonly used *16S rRNA* primer pairs using environmental DNA and cDNA from soil and sediment samples. To directly test the ecological validity of the *rpoB* transcriptional signal and determine if it reflects functional activity, the approach was applied to batch experiments from a Biological Activated Carbon (BAC) biofilter system in which dissolved organic carbon (DOC) removal rates were measured in parallel with *rpoB* and *16S rRNA* gene and transcript abundances across a time series. We demonstrate that *rpoB* captures bacterial community composition comparable to *16S rRNA* while providing transcriptional signals more strongly associated with active microbial populations. This work establishes *rpoB* as a robust activity-responsive complementary marker for environmental microbial ecology and provides a framework for activity-resolved microbiome profiling.

## 2. Materials and Methods

### 2.1 Proteins coding gene screening and candidate selection

#### 2.1.1 rpoB nucleotide database construction and curation, de-novo primer design and comparison with existing rpoB primer pairs

*rpoB* nucleotide sequences and taxonomy were collected from six sources (Supplementary Table 1) and curated using custom R scripts to retain full-length sequences and partial-length nucleotide sequences (2,000bp or longer) that contained a truncated open reading frame (ORF). This resulted in a curated database containing 305,274 (269,903 full length) unique *rpoB* sequence*s* (*rpoB* Taxonomy reference database; Supplementary Figure 1). From this a subset database of 21,763 full length, unique sequences with full taxonomic resolution to species level was made (*rpoB* Primer database; Supplementary Figure 1) for primer design and testing using Primer Prospector (Walters et al., 2011). Only primer pairs matching the criteria of this study (primers with degeneracies ≤ 384 generating amplicons suitable for both q-PCR and Illumina sequencing *i.e.* amplicon length between 200bp and 600bp, and consistent size across different species), were retained. The performance of different *rpoB* primer pairs was compared using the in-silico PCR approach described previously (Okonkwo et al., 2023; Supplementary Material and Methods).

#### 2.1.2 Comparison of rpoB to other protein-coding markers genes

The best performing *rpoB* primers were then compared with other candidate marker genes selected from a list of single copy core genes used for phylogenetic tree reconstruction (Na et al., 2018) and genes for which universal bacterial PCR primers have been proposed (Santos and Ochman 2004; Barret et al., 2015). This list was further restricted to genes available in the GTDB database (n=79; Supplementary Table 2). De-novo primers for each gene were designed with Primer Prospector (Supplementary Figure 1; Supplementary Material and methods) and these, along with any existing primers (Supplementary Table 3) matching the study criteria, were evaluated by in-silico PCR. The primer pairs with the best in silico performance (four and one *rpoB* and *gyrB* primer pairs, respectively) were tested by end-point PCR and (RT)-qPCR (Supplementary Material and methods).

### 2.2 *16S rRNA* primer selection and in silico comparison to *rpoB*

The coverage of the best performing *rpoB* primers (de-novo 1528F/2041R) was compared with the coverage of six commonly used *16S rRNA* primers (Supplementary Table 3), selected from a list of nine *16S rRNA* primers based on their in-silico PCR performance against the Ribogrove database of full-length *16S rRNA* genes (Sikolenko and Valentovich, 2022) (Supplementary Material and methods; Supplementary Figure 1). A database containing 16,922 full length nucleotide sequences from unique bacterial species with both *rpoB* and *16S rRNA* was made from our *rpoB* database and MIMt database (Cabezas et al., 2024). The weighted score (WS) of the best performing *16S rRNA* primers pair (515F/926R) and *rpoB* primer pair (1528F/20141R) were calculated separately for each individual bacterial phyla to predict differences in coverage within the bacterial kingdom.

### 2.3 *rpoB* verses *16S rRNA* for (RT)-q-PCR and (RT)-amplicon sequencing in environmental and BAC batch samples

Surface soil and river sediments (n=5) were collected in Kelvingrove Park (Glasgow, UK [55.870204, -4.291753]), September 2024 and stored at -80°C. Biological Activated Carbon (BAC) samples were obtained from a lab-scale biofilter experiment (Shi et al., 2024) and incubated in sterilised glass bottles in the presence of 0.2µm filtered surface water (∼1.5g BAC plus 50ml water) at 10°C (BAC 10) and 20°C (BAC 20) and destructively sampled after 0, 8, 24, 48 and 72 hours. Dissolved organic matter (DOC) removal rates were measured as the difference in DOC concentrations in influent water (µg DOC) at each time points compared to the previous time point, divided by the time interval (µg DOC removed/ hour). At each time point, BAC also was collected and stored at -80°C prior to DNA and RNA extraction. DNA/RNA co-extraction was performed using commercial kits (AllPrep PowerFecal Pro DNA/RNA Kit (Qiagen) and the ZymoBIOMICS DNA/RNA miniprep kit (Zymo Research), RNA integrity was evaluated using the Ramp (Cholet et al., 2019) methods and RNA reverse transcription was performed using the SuperScript IV kit (Thermo Fisher Scientific, UK). Amplicon libraries (DNA and cDNA) were prepared as in Cholet et al., (2019) and sequenced using Illumina NextSeq. ASVs were then constructed following the DADA2 pipeline and quality checked for rpoB following the recommendations from Cholet et al., (2022). (Full details in Supplementary Material and Methods).

## 3 Results

### 3.1 *rpoB* 1528F/2041R outperforms all candidate protein-coding marker gene primers for bacterial community profiling

#### 3.1.1 De novo primer design and comparison with existing rpoB pairs

De novo primer design yielded three candidate *rpoB* primer pairs meeting the study criteria (degeneracy ≤384; amplicons between 200–600 bp with consistent size): 1528F/2041R (∼533 bp), 3304F/3817R (∼533 bp), and 3304F/3826R (∼542 bp). The 1528F/2041R pair had the lowest degeneracy of all candidate pairs (12 and 24 for forward and reverse primers, respectively). In silico PCR showed that 1528F/2041R (29% at WS=0; 77% at WS=1; 97% at WS=3) matched or outperformed all other de novo and published *rpoB* primers suitable for amplicon sequencing, including the modified Ogier et al., (2019) pair (29%; 73%; 96%) and Leclerc et al., (2025) (29%; 77%; 96%). The highest-performing existing primer set, Bivand et al., (2024) (∼67% at WS=0; ∼96% at WS=1; ∼98% at WS=3) (Supplementary Table 3) achieved marginally superior coverage at low mismatch thresholds but at a cost of substantially higher degeneracy (up to 12,000 and 98,000 for the forward and reverse primers, respectively), which is prohibitive for qPCR and amplicon sequencing applications. The 3304F/3817R and 3304F/3826R pairs were excluded because they predicted amplicons of multiple lengths (Supplementary Figure 2) and Leclerc et al., 2025 pair was not considered due to amplicon length (∼907bp). Three primer pairs Bivand et al., 2024, 1528F/2041R (this study) and modified Ogier et al., 2019 set (this study) were selected for progression based on coverage and amplicon size.

#### 3.1.2 Comparison with primers targeting other protein-coding marker genes

The three best performing *rpoB* primers were benchmarked against 79 other protein-coding marker genes. Out of these 79 genes, de-novo primers could be found for 33 genes (Supplementary data 1; Supplementary Table 2) with only seven of these matching our criteria (Supplementary Table 3). The best de novo alternative was the *atpD* 636F/836R pair (∼12% at WS=0; ∼75% at WS=1; ∼96% at WS=3), which performed comparably to the modified Ogier et al., (2019) *rpoB* pair but fell below the 1528F/2041R and Bivand et al., (2024) pairs at all thresholds (Figure 1B). Among previously published multi-gene primer sets, only the *gyrB* primer pair of Barret et al., (2015) achieved coverage approaching the best *rpoB* primers (∼21% at WS=0; ∼66% at WS=1; ∼94% at WS=3) with consistent amplicon length. All other published protein-coding marker gene primers (targeting *ileS*, *lepA*, *leuS*, *pyrG*, *recA*, *recG*, *rplB*; Santos & Ochman, 2004) showed lower coverage at every WS threshold (Figure 1C). These results identify *gryB* and *rpoB* as the protein-coding marker genes with the broadest bacterial coverage suitable for combined qPCR and amplicon sequencing.

**Figure 1.**
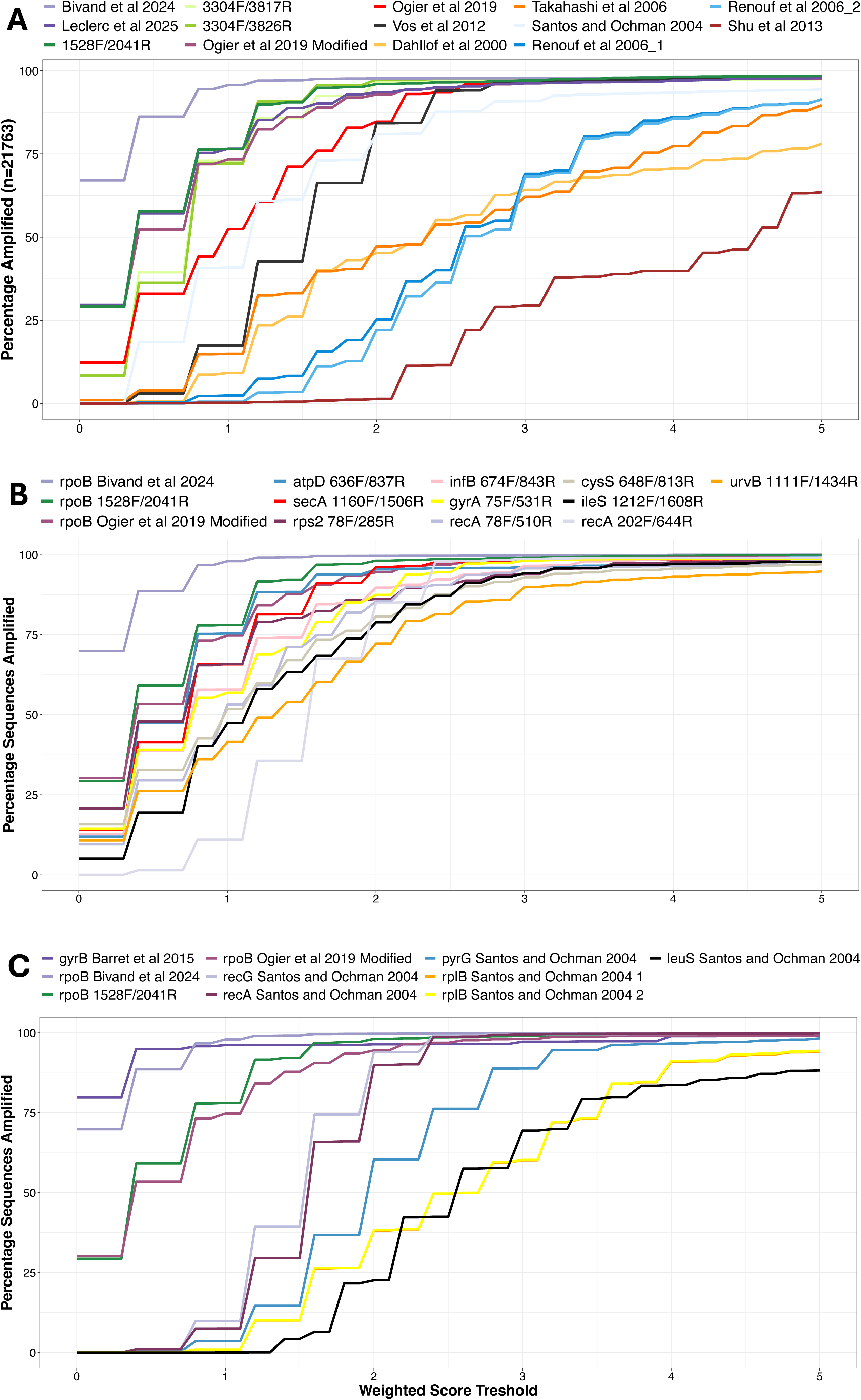
In-silico PCR comparison of coverage for primer pairs targeting protein-coding marker genes. The best performing de-novo and existing *rpoB* PCR primer pairs (A) were compared to de-novo (B) and exiting (C) primer pairs targeting other protein-coding marker genes, considering WS thresholds between 0 and 5.

#### 3.1.3 Experimental PCR and qPCR validation

The three best-performing *rpoB* primer pairs and the *gyrB* Barret et al., (2015) pair were taken forward for experimental validation using soil DNA and cDNA. Only the 1528F/2041R and modified Ogier et al., (2019) pairs produced clear, correctly-sized amplicons after 25 PCR cycles from both soil DNA and cDNA, consistent with in silico predictions; the remaining *rpoB* pairs and the *gyrB* pair required 35 cycles and yielded weak or ambiguous bands and were excluded from further analysis (Supplementary Figure 4). Between the two retained pairs, 1528F/2041R produced consistently stronger cDNA amplification. The 1528F/2041R pair was therefore selected for qPCR optimisation, yielding an optimal annealing temperature of 54.5°C (non-barcoded) and 60°C (barcoded), with qPCR efficiencies of ∼81% (DNA standard) and ∼90% (RNA standard) (Table 1; Supplementary Figure 5). Taken together, across in silico coverage, primer degeneracy, amplicon uniformity, and experimental performance, the de novo 1528F/2041R pair is identified as the best-performing *rpoB* primer set for both qPCR quantification and amplicon sequencing.

**Table 1.**
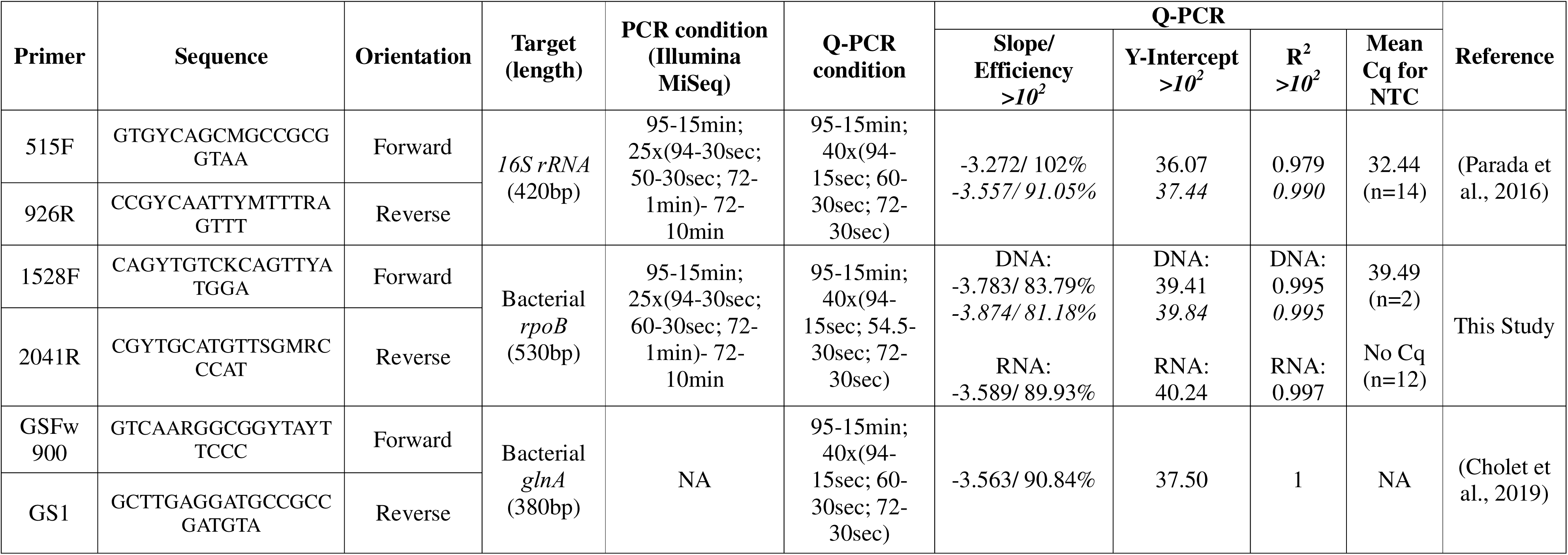
List of Primers and PCR conditions used in this study.

### 3.2 *rpoB* 1528F/2041R provides robust predicted coverage of dominant bacterial phyla despite lower absolute breadth than *16S rRNA*

To contextualise the coverage of the new 1528F/2041R *rpoB* primers, they were compared in silico with six widely used *16S rRNA* primer pairs (V2f/V3r, 341F/785R, 341F/806R, 515F/806R, 515F/926R and 969F/1406R) using a matched database of 16,922 bacterial species present in both the *rpoB* and MIMt *16S rRNA* databases. Primer pairs 27F/338R, 63F/338R and 1369F/1389P/1492R were excluded because of partial-length sequences in the MIMt database and lower in silico performance against the Ribogrove database (Sikolenko and Valentovich, 2022). Five of the six *16S rRNA* primer pairs amplified ∼95% of species at WS=0, with 515F/926R performing best across all thresholds and selected as the benchmark. The *rpoB* 1528F/2041R pair amplified ∼30% of species at WS=0, increasing to ∼78% at WS=1 and ∼97% at WS=3 (Figure 2A). Although coverage was lower at strict mismatch thresholds, the gap narrowed substantially at WS≥1, which is more representative of environmental amplicon studies. At the phylum level, *rpoB* 1528F/2041R showed good coverage (average combined WS≤1) for 29 of 42 bacterial phyla, including Actinomycetota (WS=0.22; n=3,751), Pseudomonadota (WS=0.32; n=6,211), Bacteroidota (WS=0.68; n=2,196), and Bacillota (WS=0.71; n=3,143), which together represented ∼90% of all sequences. Coverage was intermediate (WS 1-2) for ten phyla, while Campylobacterota (WS=2.8), Coprothermobacterota (WS=3.2), and Planctomycetota (WS=3.3) showed poor coverage, largely due to the forward primer 1528F (average WS=3.34 versus 0.22 for 2041R). In comparison, the *16S rRNA* 515F/926R pair achieved WS≤1 across all 42 phyla and WS≤0.1 for 39 phyla (Figure 2B).

**Figure 2.**
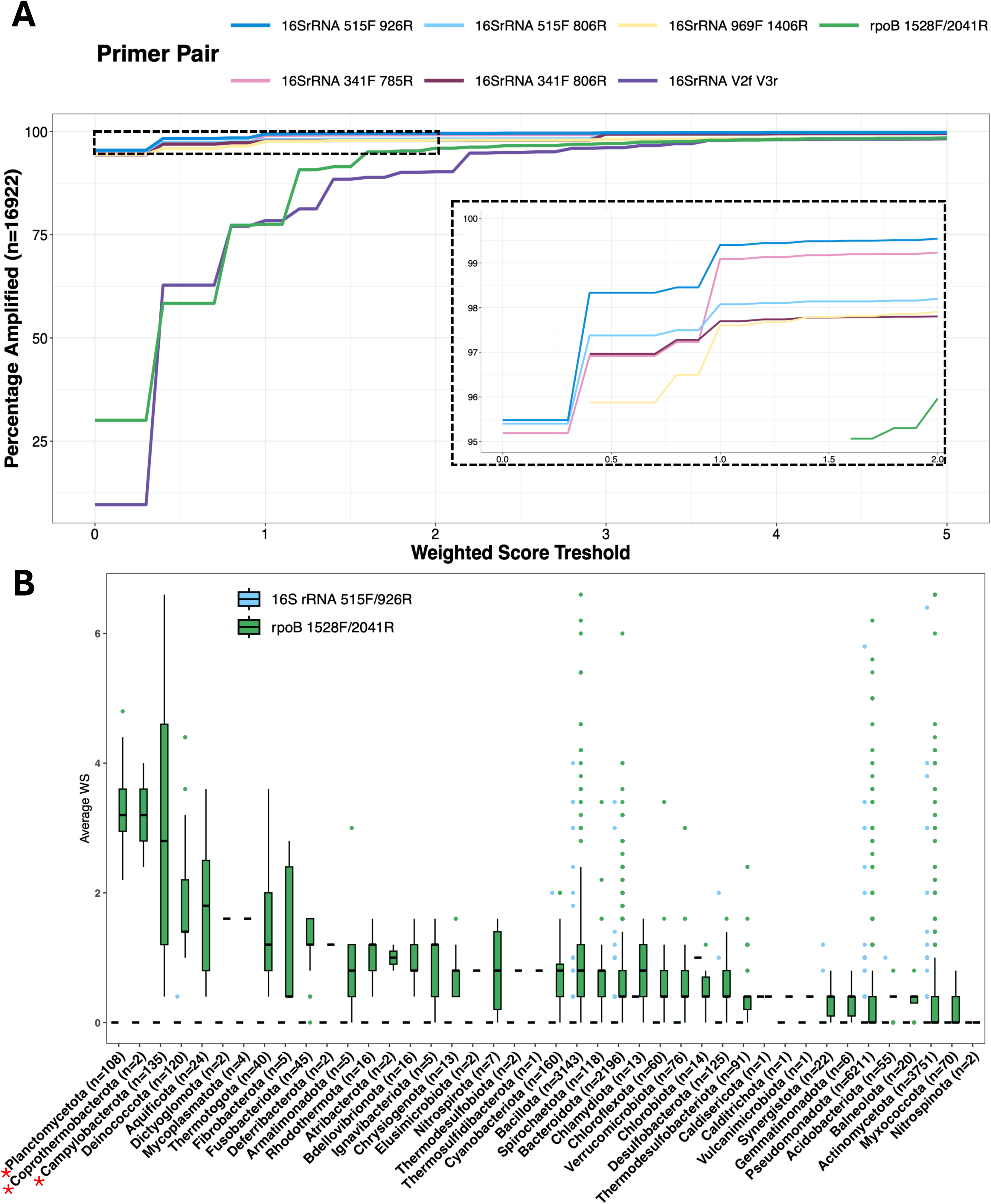
In-silico comparison of *rpoB* and *16S rRNA* primer pairs. In-silico PCR were carried out for the best performing *rpoB* primer pair and commonly used *16S rRNA* primer pairs against overlap *rpoB* database and overlap MIMt database, respectively (A). The combined average WS of forward and reverse primers for the *16S rRNA* primer pair 515F/926R and *rpoB* primers 1528F/2041R for each bacterial phylum is presented in (B). Phyla with average combined WS >2 are shown by red stars *.

Overall, while *16S rRNA* primers retained greater taxonomic breadth, *rpoB* 1528F/2041R provided robust predicted coverage of the dominant bacterial lineages in environmental samples. We next tested whether it could offer the additional advantage of capturing transcriptional activity.

### 3.3 *rpoB* qPCR assay achieves two orders of magnitude greater sensitivity and lower background than *16S rRNA*

A direct comparison of qPCR performance revealed markedly greater sensitivity for *rpoB* than *16S rRNA*. The *rpoB* 1528F/2041R assay had a limit of detection (LoD) of 8 copies/µl, defined as the concentration yielding a 95% detection probability (1 of 6 replicates failed at 1.6 copies/µl), and a limit of quantification (LoQ) of 8-40 copies/µl. Amplification in no-template controls (NTCs) was rare, occurring in only 2 of 14 reactions (mean Cq = 39.49). In contrast, a *16S rRNA* LoD could not be established because all 14 NTCs amplified (mean Cq = 32.35). Its LoQ was 1,000-3,200 copies/µl, approximately two orders of magnitude higher than *rpoB*, due to loss of linearity below 3,200 copies/µl. These results demonstrate a substantially greater quantification range and sensitivity for *rpoB* qPCR.

In environmental samples, *16S rRNA* gene copy numbers were 5-7-fold higher than *rpoB* in both soil (3.53 × 10□ vs. 6.68 × 10□ copies g□¹ wet weight) and sediment (3.01 × 10□ vs. 4.54 × 10□ copies g□¹), consistent with the multicopy nature of the ribosomal gene. Transcript patterns differed markedly between markers. *16S rRNA* transcripts exceeded gene copies by ∼300-fold in soil and ∼1,300-fold in sediment, whereas *rpoB* transcripts were ∼190-fold and ∼160-fold lower than gene copies, respectively. This contrast was unlikely to result from RNA degradation, as high Ramp values (0.86-0.90) indicated excellent mRNA integrity (Cholet et al., 2019), and the *glnA* cDNA:DNA ratio showed a similar pattern to *rpoB* (∼170-fold and ∼50-fold lower than gene copies in soil and sediment, respectively; Supplementary Figure 6).

### 3.4 *rpoB* and *16S rRNA* recover concordant dominant communities but diverge in rare taxa detection and transcriptional resolution

*3.4.1 Comparison of rpoB and 16S rRNA DNA community profiles*.

*rpoB* amplicon sequencing using single-end reads retained more reads than the best paired-end approach (∼94% and ∼88% in soil DNA and cDNA; ∼94% and ∼83% in sediment versus ∼83% and ∼71% in soil and ∼83% and ∼60% in sediment), with also higher proportions of sequences translating to a correct protein (∼99% vs. ∼96%) and fewer unassigned ASVs (Supplementary Table 4). All subsequent analyses therefore used the single-end approach. At the phylum level, *rpoB* detected 21 phyla in soil and 22 in sediment, compared with 26 and 32 for *16S rRNA*. Most phyla were shared (18 across both environments) and accounted for >85% of community abundance, whereas phyla unique to either marker each represented <0.5% of total abundance, indicating that primer-related differences were largely confined to the rare biosphere (Figure 3).

**Figure 3.**
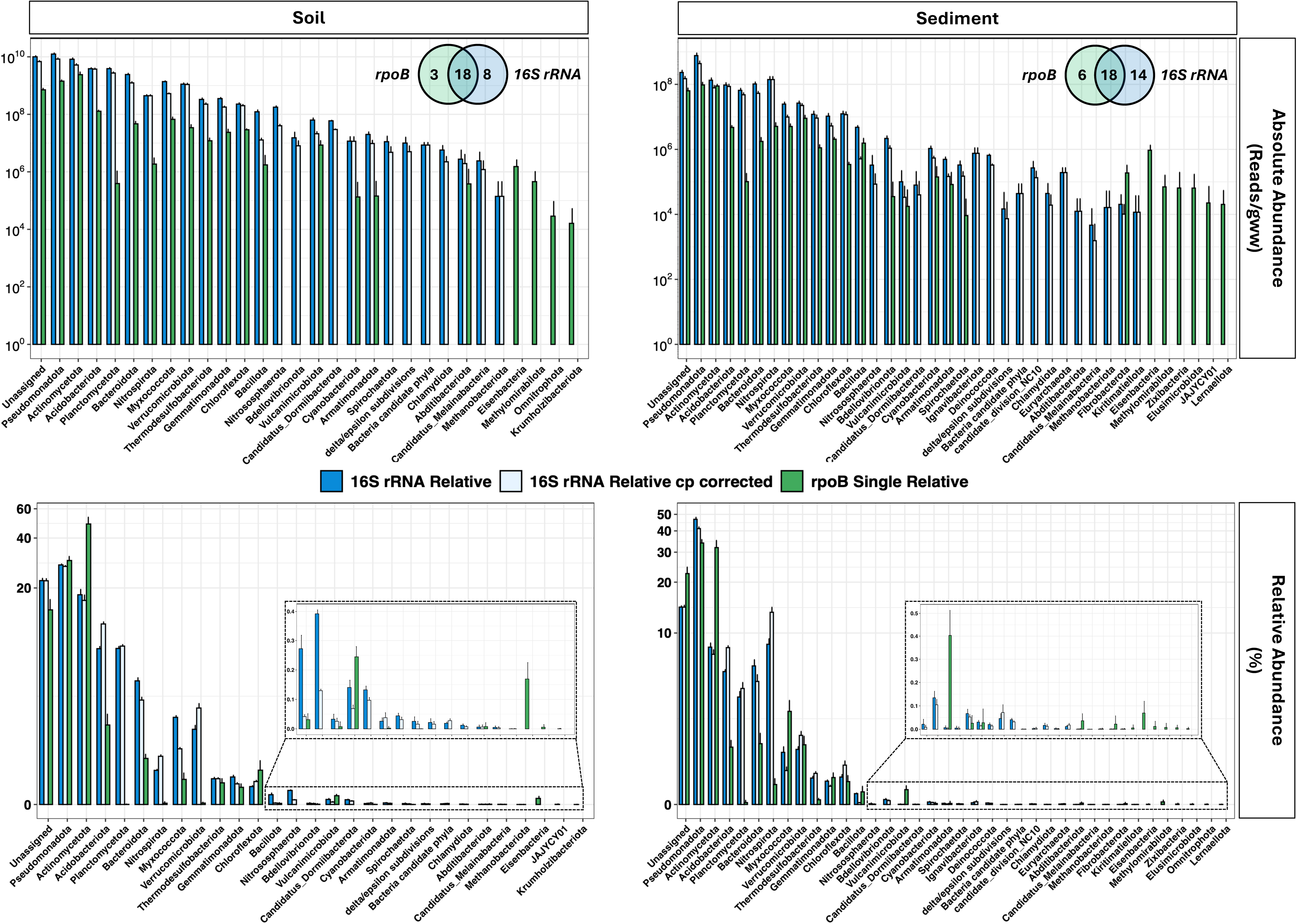
Abundance of bacterial phyla in soil (left) and sediment (right) based on absolute (top) and relative (bottom) values. Phyla abundances are presented as absolute (Top) or relative (bottom) values. For *16S rRNA*, abundances are presented with (*16S rRNA* Normalised) or without (*16S rRNA*) correction for estimated copy numbers. The Venn Diagrams on absolute abundance plots represent the number of phyla shared between *16Sr RNA* and *rpoB* datasets.

Phyla unique to *16S rRNA* included Archaeal lineages (Nitrososphaerota, Methanobacteriota, and in sediment Euryarchaeota), and candidate phyla (Candidatus Dormiibacterota, candidate_division_NC10 and Candidatus Melainabacteria), expected, as the *rpoB* primer was designed for Bacteria. Planctomycetota was more abundant in *16S rRNA* data, consistent with the poor predicted coverage of *rpoB* primer 1557F. Conversely, *rpoB* uniquely detected several low-abundance bacterial lineages, including Eisenbacteria and Krumholzibacteriota (soil and sediment), and Methylomirabilota, Zixibacteria, Elusimicrobiota, Omnitrophota and Lernaellota (sediment). Thus, both markers recovered the dominant taxa, while each detected distinct rare lineages, supporting *rpoB* as a complementary rather than replacement marker for *16S rRNA*. Absolute phylum-level abundances, estimated from qPCR (and estimated copy-number for *16S rRNA*) normalised data were significantly higher for *16S rRNA* than *rpoB* in both soil and sediment (Figure 3). However, this was not due to greater taxonomic richness. Within the four dominant shared phyla, *rpoB* consistently recovered more unique taxa at the Order, Family, and Genus level for Acidobacteriota, Actinomycetota, and Pseudomonadota in both soil and sediment, except for Pseudomonadota (Family) in both environments and Order level in soil. (Figure 4).

**Figure 4.**
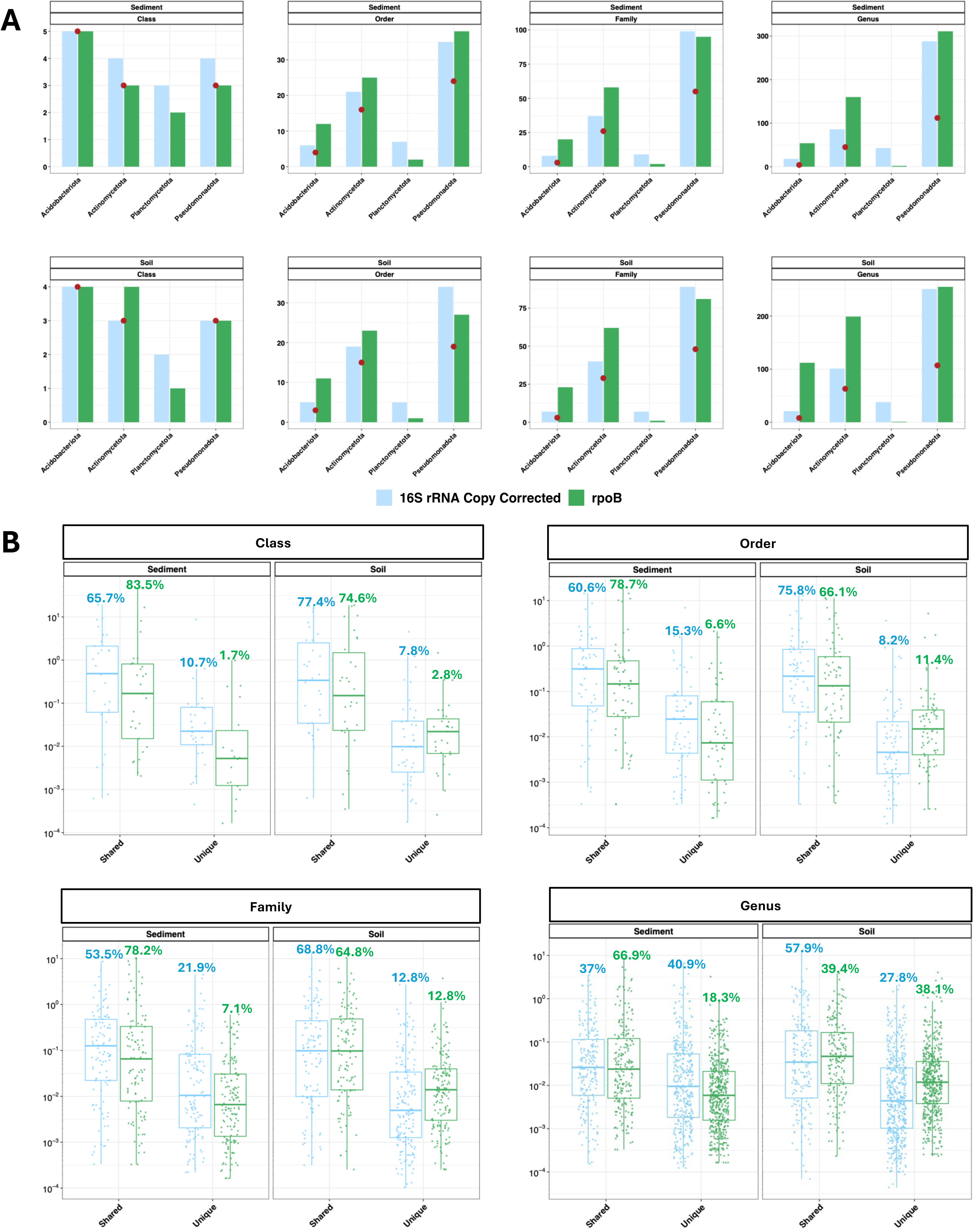
Distribution of shared and unique richness between *16S rRNA* and *rpoB*. (A) Richness within the four most abundant phyla at different phylogenetic levels from *16S rRNA* and *rpoB* sequencing of the same soil and sediments. Red dots indicate the number of shared taxa between gene targets. (B) The percentage of shared and unique taxa between the *16S rRNA* and *rpoB* sequencing datasets (relative to their respective total community) at different phylogenetic levels in soils and sediments

*3.4.2 Comparison of rpoB and 16S rRNA DNA and cDNA community profiles*

Both markers detected more ASVs in cDNA libraries than in corresponding DNA libraries, indicating apparent greater diversity in the transcriptionally active fraction. However, the patterns differed substantially between markers. For *16S rRNA*, ∼50% of DNA ASVs were also found in cDNA libraries (53.5% soil; 48% sediment); for *rpoB* the overlap was lower (∼30%; 31.4% soil, 27.7% sediment), suggesting greater divergence between the total and active communities as captured by *rpoB*. In both cases, the shared ASVs accounted for the vast majority of sequence abundance (>85% for *16S rRNA*; ∼60–70% for *rpoB*), confirming that unique ASVs were predominantly rare taxa (Table 2).

**Table 2.**
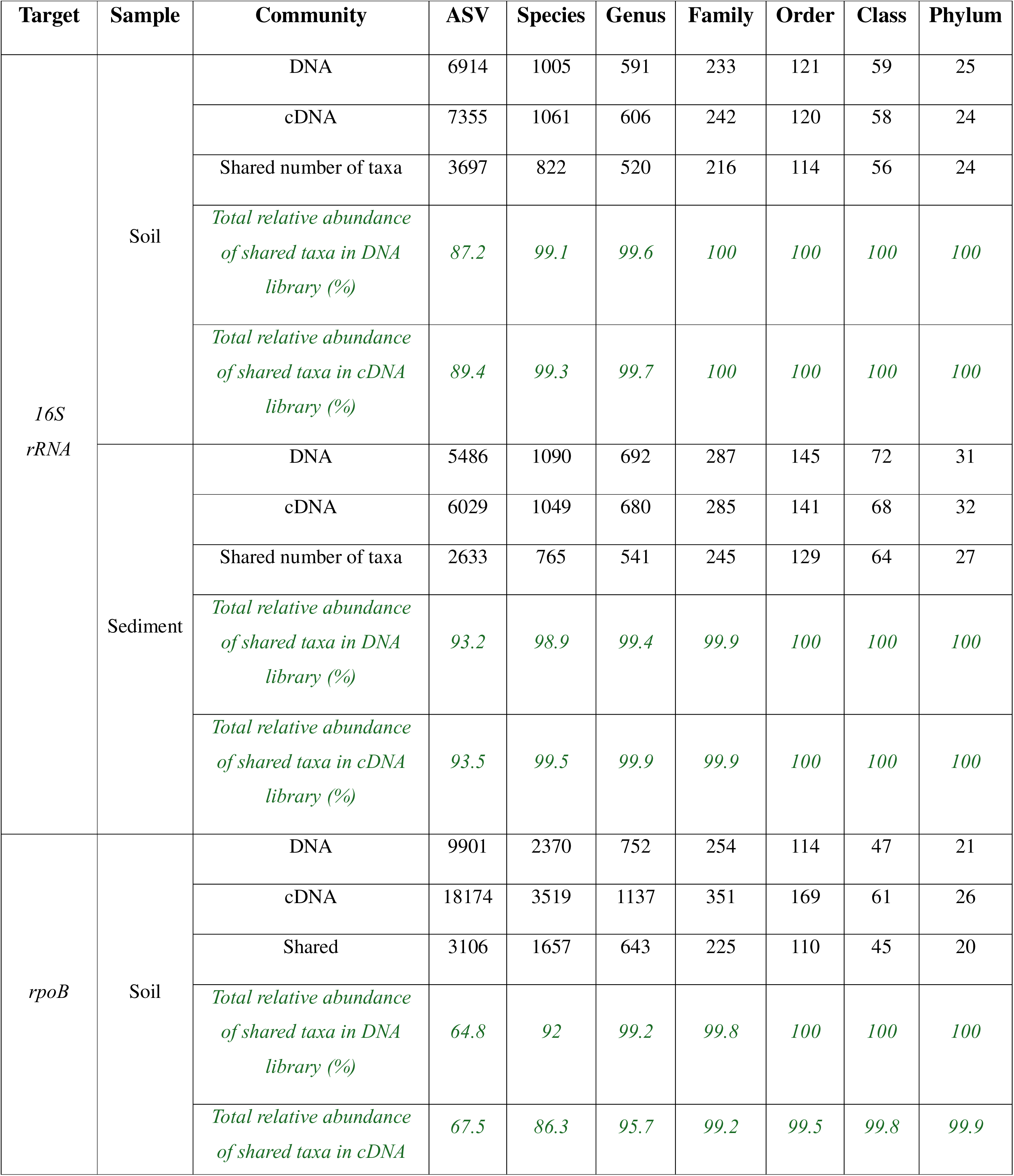
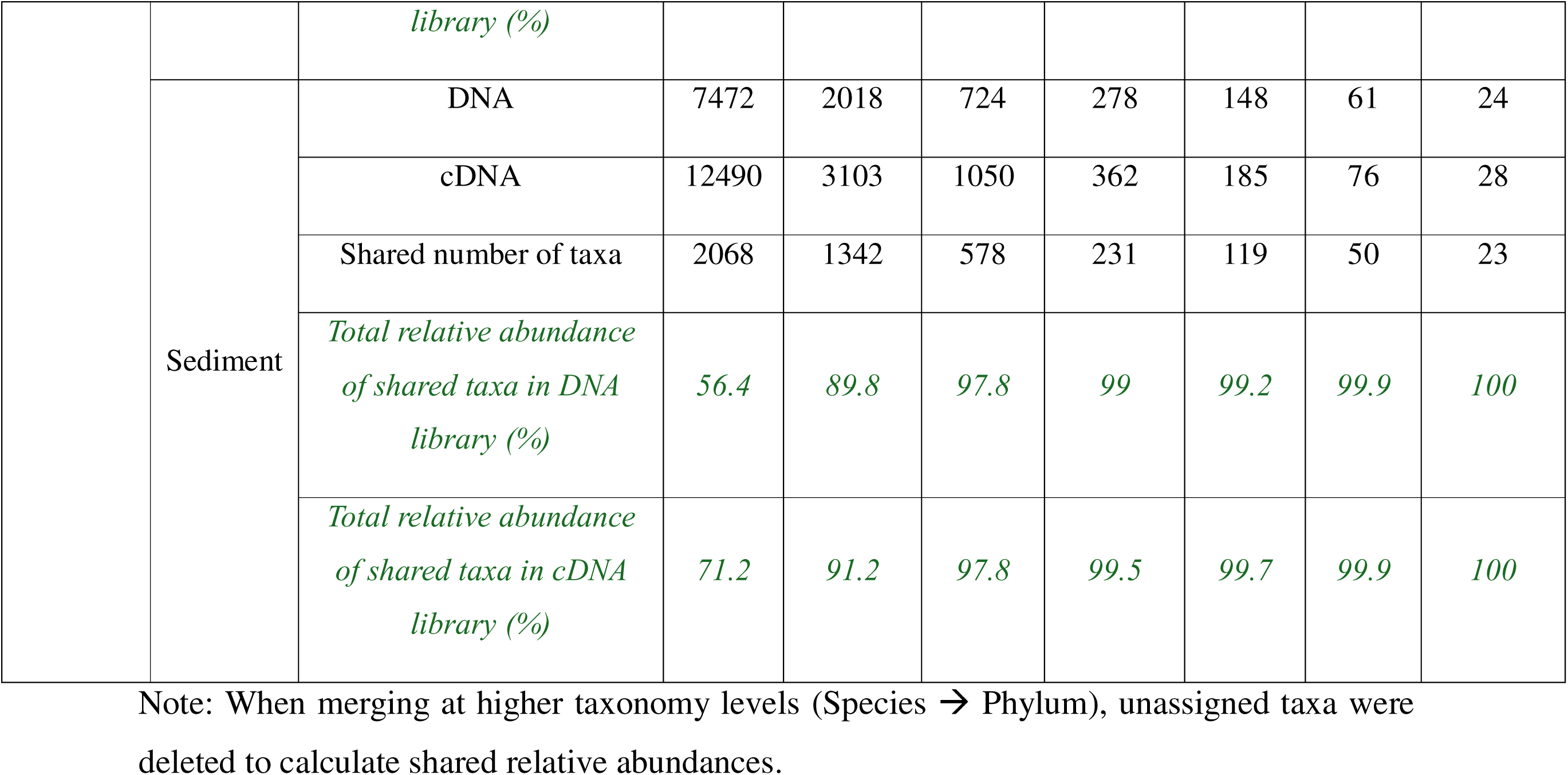
Richness of total (DNA) and transcriptionally active (cDNA) communities from ASV to phylum level in soil and sediment revealed to 16Sr RNA and rpoB sequencing.

The most transcriptionally active genera identified by absolute transcript abundance showed broad concordance between markers: eight genera in soil and ten in sediment were shared between the *16S rRNA* and *rpoB* cDNA datasets, including *Bradyrhizobium* and *Hyphomicrobium* in both soil and sediment. Limited overlap at genus level in some cases likely reflects database limitations rather than true biological differences, as active genera identified by *16S rRNA* were often recovered by *rpoB* at the family or order level (e.g. Vicinamibacterales, Haliangiaceae) (Figure 5A–D).

**Figure 5.**
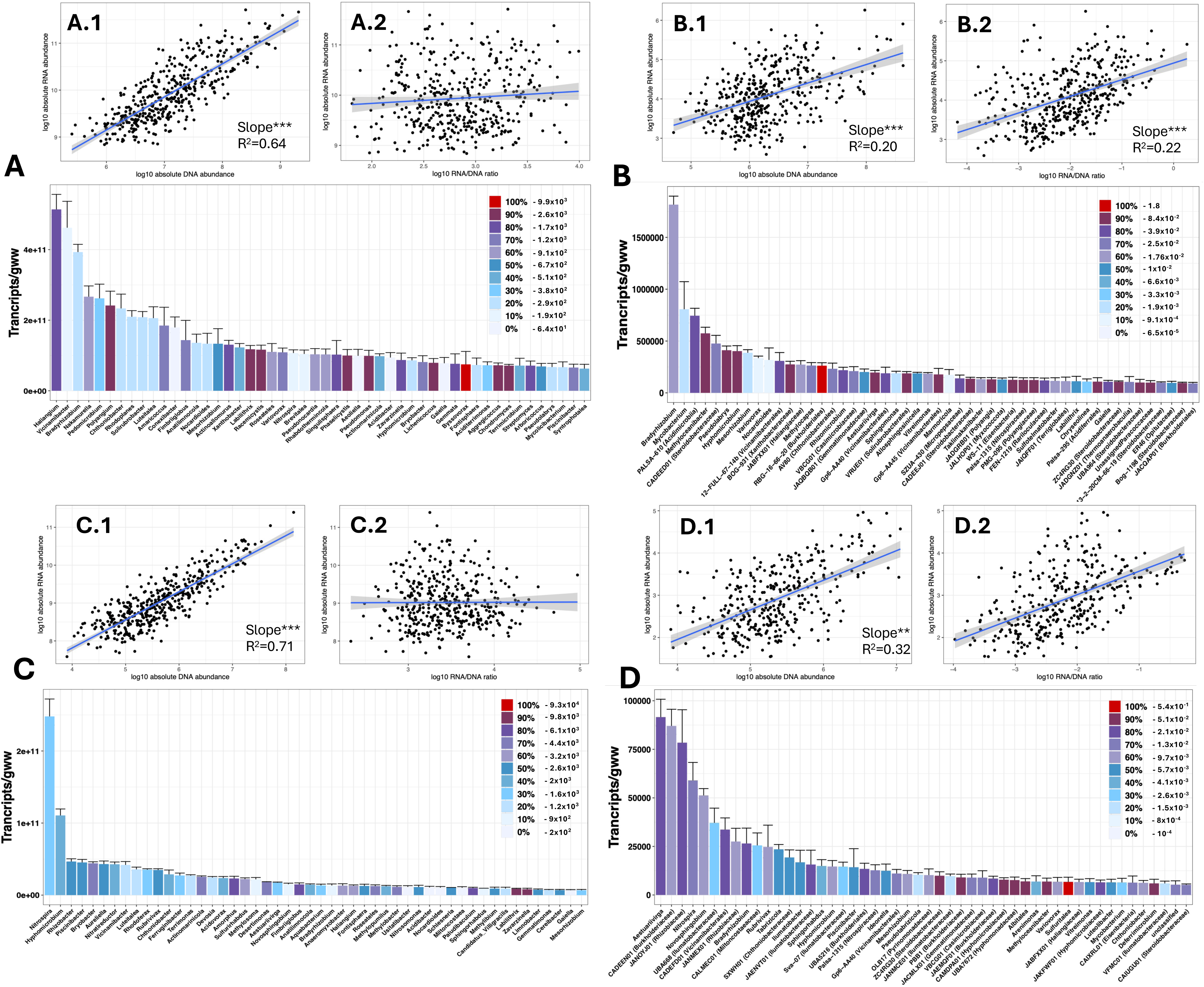
Transcriptionally active communities in soil and sediment. Y-axis show the absolute transcript abundance of bacterial genera (top 50 most transcriptionally active) in soil for *16S rRNA* in soil (A) and *rpoB* (B) and sediment for *16S rRNA* in soil (C) and *rpoB* (D). The colour of the bars indicate the RNA/DNA ratio quantile of individual taxa as indicated in the legend (numbers associated to quantile are added to individual legends). The regressions between absolute cDNA and absolute DNA abundance (log10) and between absolute cDNA abundance and between cDNA/DNA ratio (log 10) are shown are shown in A.1/B.1/C.1/D.1 and A.2/B.2/C.2/D.2, respectively.

### 3.5 *rpoB* cDNA:DNA ratios track taxon-level transcriptional activity where *16S rRNA* ratios do not

The most important finding concerns the relationship between transcript abundance and the cDNA:DNA ratio, a key metric for assessing transcriptional activity at the taxon level. For *16S rRNA*, cDNA:DNA ratios ranged from 10²–10³ (lowest) to 10³–10□ (highest) across genera. There was no significant relationship with absolute transcript abundance (slope p > 0.5 in both soil and sediment; Figure 5A.2, B.2). For *rpoB*, cDNA:DNA ratios spanned a wider dynamic range, from 10□□ to ∼1.8, and exhibited a significant positive correlation with absolute transcript abundance in both soil (R²=0.22, p<0.001) and sediment (R²=0.29, p<0.001) (Figure 5C.2, D.2). This relationship held across environments and was not driven solely by co-variation between gene and transcript copy numbers: while DNA and cDNA abundances were strongly correlated for *16S rRNA* (R²=0.64 soil; R²=0.71 sediment; p<0.001), the equivalent relationship for *rpoB* was weaker (R²=0.20 soil; R²=0.32 sediment), indicating that *rpoB* transcript levels reflect transcriptional state rather than simply tracking gene copy number.

Comparing□*rpoB*□and□*16S*L*rRNA*□transcript data shows that while both markers capture broadly similar active taxa, *rpoB* revealed stronger differentiation among actively transcribing taxa and a measurable, positive link between transcript abundance and cDNA:DNA ratios.□Whereas□1*6S*L*rRNA*□transcripts largely mirror total biomass,□*rpoB*□captures taxa-specific transcriptional dynamics.

### 3.6 *rpoB,* but not *16S rRNA* transcripts, quantitatively tracks changes in carbon degradation rates

In batch series incubations, changes in DOC removal rates, operated and incubated at 10 and 20°C (BAC10 and BAC20), were followed alongside changes in *16S rRNA* and *rpoB* gene and transcript abundances. The DOC removal rates decreased through time for both BAC10 (∼268µg/h, ∼41µg/h, ∼28µg/h and ∼11µg/h for the intervals 0h-8h, 8h-24h, 24h-48h-72h, respectively) and BAC20 samples (∼295µg/h, ∼84µg/h, ∼29µg/h and ∼17µg/h for the intervals 0h-8h, 8h-24h, 24h-48h-72h, respectively) (Figure 6A). This downward trend was matched by the decrease in *rpoB* transcript copy number through time for both BAC10 (-44%, -86%, -97% and -98% at time 8h, 24h, 48h and 72h compared to time 0, respectively) and BAC20 (-65%, -98%, -99% and -99% at time 8h, 24h, 48h and 72h compared to time 0, respectively), while *rpoB* gene copy numbers remained relatively constant at all time points (Figure 6B). Both the *rpoB* transcripts abundances and transcription ratios (cDNA/DNA) correlated well (correlation coefficients between 0.84 and 0.99) with the DOC removal rates (Supplementary Table 7). Comparatively, little change in *16S rRNA* transcripts were observed, with fluctuations through time for BAC10 (+31%%, -9%, +25% and -9% at time 8h, 24h, 48h and 72h compared to time 0, respectively) and a slight downward trend for BAC20 (+48%%, -27%, -36% and -28% at time 8h, 24h, 48h and 72h compared to time 0, respectively) while *16S rRNA* gene copy numbers also remained constant through time (Figure 6B). Consequently, *16S rRNA* transcripts abundances and transcription ratios (cDNA/DNA) showed inconsistent correlation with DOC removal rates (from -0.4 to 0.98) (Supplementary Table 7).

**Figure 6.**
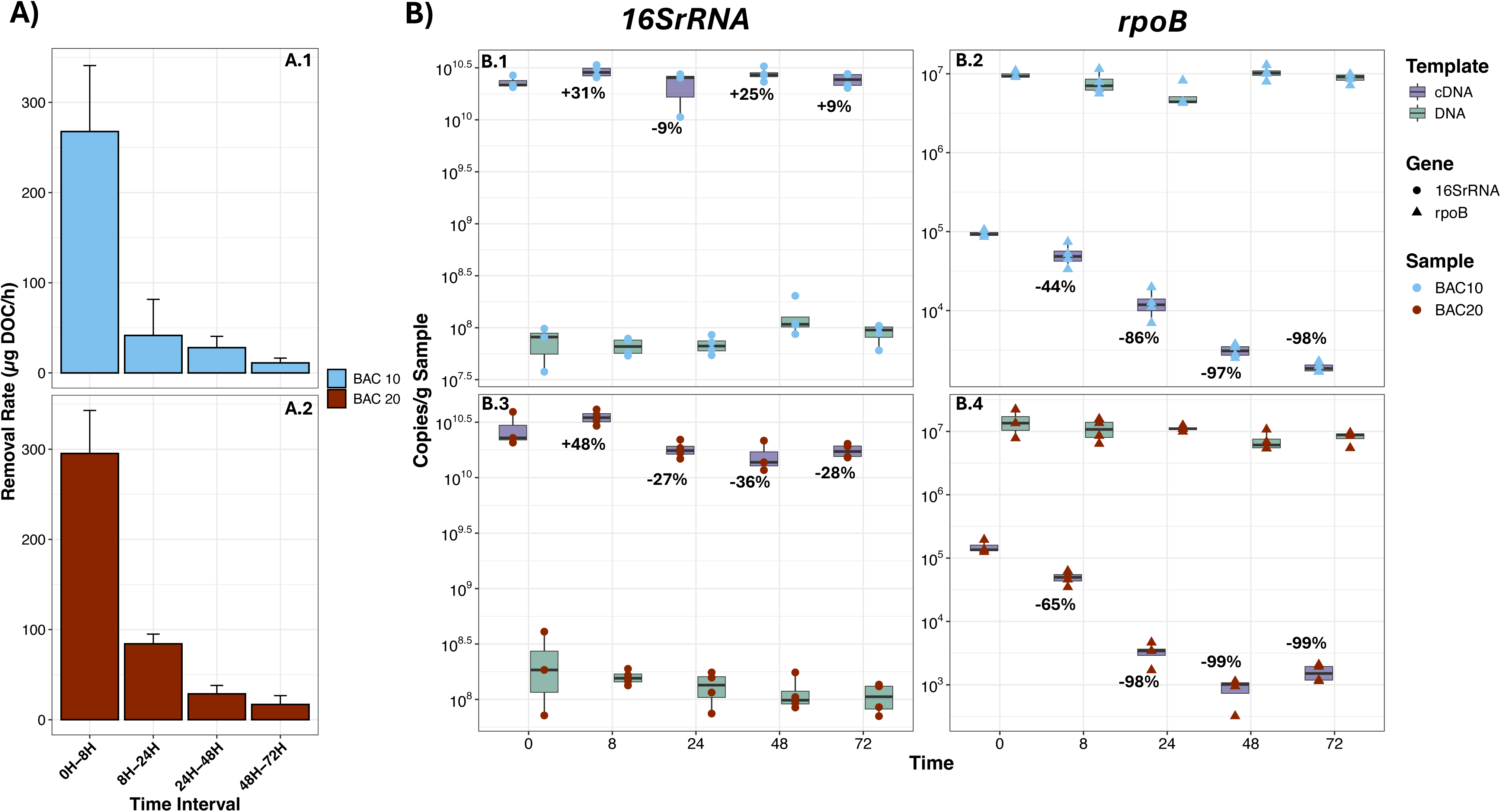
Relationship between changes in transcriptional activity and process rate. A: Changes in DOC removal rates (µg/hour) between successive time points. B: Changes in gene (DNA) and transcript (cDNA) for *16S rRNA* (B.1 and B.3) and *rpoB* (B.2 and B.4); Percentage changes in transcript abundance were calculated compared to Time 0.

## 4. Discussion

### 4.1 Combining *rpoB* DNA and cDNA Q-PCR and sequencing to identify transcriptionally active bacteria in natural environment

This study provides the first systematic evaluation of protein-coding marker genes, including target selection, primer design and experimental validation on DNA and cDNA templates to identify complements of *16S rRNA* for bacterial taxonomic composition and transcriptional activity in environmental samples. By benchmarking the new *rpoB* 1528F/2041R primers against the field-standard *16S rRNA* approach, we show that alternative markers can complement ribosomal profiling and potentially capture metabolically active populations more accurately. The finding that *rpoB* cDNA:DNA ratios scale with transcript abundance across taxa, whereas *16S rRNA* ratios do not (Figure 5), and that *rpoB* transcript copy numbers better reflect changes in process rates (Figure 6), represent the central results of this study. These findings suggest that *rpoB* expression captures taxon-level variation in transcriptional activity, a capability that *16S rRNA*-based approaches cannot readily provide. Barnard et al. (2013, 2015) previously showed that increases in process rates were accompanied by higher *rpoB* transcript abundances while gene abundances remained constant, indicating elevated transcript numbers per cell in active bacteria, whereas *16S rRNA* transcripts did not track these changes. Here, we further show that *rpoB* transcript abundance and cDNA:DNA ratios decline with decreasing process rates, while *rpoB* gene copy numbers and, importantly, *16S rRNA* transcript abundances remain relatively constant. Previous studies have also shown that *rpoB* transcripts respond to stress-induced shifts in metabolism (Peng et al., 2017), and reflect activity associated with pollutant degradation (Gofstein and Leigh, 2023). Together, these studies and our results indicate that changes in *rpoB* transcript abundance can serve as quantitative indicators of bacterial metabolic state, providing a scalable alternative to metatranscriptomics for assessing transcriptional activity in complex bacterial communities. Across all samples, *rpoB* transcript levels were consistently lower than gene copy numbers (∼150 fold in sediment and ∼180 fold in sediment). High RNA integrity, confirmed by Ramp values, together with agreement between *rpoB* and *glnA* cDNA:DNA ratios, indicates that these low ratios are not artefacts of RNA degradation or assay inefficiency. Barnard et al., (2013, 2015) similarly reported *rpoB* transcript abundances 10³-10□-fold lower than gene copy numbers in soil, indicating overall low transcriptional activity. This interpretation is consistent with the prevalence of microbial dormancy and low metabolic activity in natural environments (Blagodatskaya & Kuzyakov, 2013; Buerger et al., 2012; Jones & Lennon, 2010; Lennon & Jones, 2011; Stenström et al., 2006).

The *16S rRNA* cDNA:DNA ratio has previously been shown to be a poor predictor of microbial growth and RNA synthesis (Papp et al., 2018). This likely reflects the complex relationship between rRNA abundance and cellular activity. Ribosome content varies with growth conditions and differs among taxa (Blazewicz et al., 2013), while intracellular rRNA decay after growth cessation depends on nutrient limitation, with rapid decay under phosphorus limitation (Himeoka et al., 2022) but accumulation of inactive rRNA during carbon starvation (Hsin-Jun Li et al., 2018). Consistent with these observations, we observed an initial increase in *16S rRNA* transcripts between 0 and 8 h, likely reflecting ribosome accumulation during the early phase of intense metabolic activity. As carbon availability declined, these ribosomes may have persisted in inactive cells, weakening the relationship between *16S rRNA* transcript abundance and DOC removal rates. This interpretation is supported by evidence that inactive and dormant cells can maintain substantial ribosome pools (Helena-Bueno et al., 2024a,b). Combined with evidence that rRNA decay can vary significantly among closely related bacterial strains (Suttner et al., 2021), these findings further confirm that *16S rRNA* transcripts are not reliable proxies for identifying active taxa within complex microbial communities.

### 4.2 *rpoB* extends community resolution while preserving ecological breadth

To facilitate broader adoption of *rpoB* in microbial ecology, we present a comprehensive synthesis and evaluation of available *rpoB* primer sets. The *rpoB* marker was compared with 79 other protein-coding genes based on conserved regions for primer design, in silico bacterial coverage, and, for selected primer sets, PCR performance. The new *rpoB* primer pair (1528F/2041R) outperformed existing and de novo primers targeting *rpoB* and other protein-coding markers in both in silico analyses and PCR-based quantification and sequencing of environmental genes and transcripts. We also describe a newly curated *rpoB* reference database containing 305,274 full-length or near-full-length unique sequences with verified protein translation, representing 20,132 additional sequences compared with GTDB. The database is freely available (Supplementary Data 3) and can be expanded as adoption of *rpoB* increases. Using this improved primer pair and database alongside the best-performing *16S rRNA* primer pair (515F/926R) and the MIMt database, we show that both approaches recovered a broad range of bacterial phyla and consistently detected the dominant lineages in soil and sediment. This agreement suggests that both markers provide comparable assessments of abundant community members. Differences were mainly confined to low-abundance taxa, with several groups uniquely detected by one marker or the other, indicating that each captures distinct components of rare biosphere diversity (Figures 3 and 4). In addition, *rpoB* often recovered greater richness within dominant phyla (Figure 4). Similar patterns have been reported in comparisons between *16S rRNA* and *rpoB* (Shu and Jiao, 2013; Vos et al., 2012) and *cpn60* (Schellenberg et al., 2009), where dominant taxa were shared but rare taxa differed between markers. Although shotgun metagenomics provides superior detection of rare taxa (Durazzi et al., 2021; Tessler et al., 2017; Durand et al., 2025), our results show that *rpoB* can reveal taxa missed by *16S rRNA* (and vice versa), supporting its use as a complementary rather than replacement marker. A key advantage of *rpoB* over *16S rRNA* is quantitative accuracy. As a single-copy bacterial gene, *rpoB* enables direct estimation of cell numbers and simplifies integration of qPCR and amplicon-sequencing data without copy number corrections. In addition, *rpoB* qPCR showed substantially lower limits of detection and quantification, making it particularly suitable for low-biomass samples. Because *rpoB* is absent from eukaryotic nuclear genomes, it is also advantageous in host-associated microbiome studies where *16S rRNA* primers can co-amplify host DNA (Stewart et al., 2024). This benefit may be especially important for low-biomass host-associated samples. Plant microbiomes may represent an exception because chloroplasts also contain a *rpoB* gene. Future improvements could include optimisation of primer coverage for Planctomycetota, potentially through the use of multiple forward primers in a single PCR reaction (e.g. Bivand et al., 2024). Probe-capture approaches applied after nucleic-acid extraction (Hiraoka et al., 2025) may also enhance detection of low-abundance genes and transcripts

Based on these results, we offer the following practical guidance for the use of *rpoB* as a marker gene: *rpoB* and *16S rRNA* could be used in combination when comprehensive detection of both abundant and rare taxa, including Archaea, Planctomycetota, and candidate phyla, is required. Where quantification of low-biomass samples is the primary goal, *rpoB* qPCR is preferred given its substantially lower detection and quantification limit, and presence as a single copy gene. For host-microbiome studies *rpoB* offers the advantage of only targeting bacteria. For studies seeking to identify taxon-level changes in transcriptional activity in response to environmental conditions or process dynamics, combined *rpoB* DNA and cDNA analysis is recommended.

## 5. Conclusion

Using integrated□in-silico, qPCR, and sequencing analyses, this study identifies□*rpoB*□and the new□1528F/2041R□primer pair as a reliable complement to□*16S*L*rRNA*□for quantifying and characterising total and transcriptionally active bacterial communities. Although its taxonomic coverage remains slightly narrower than that of□*16S*L*rRNA*,□*rpoB*□captures the dominant diversity and reveals additional low-abundance taxa while enabling measurement of□cDNA:DNA□ratios as a proxy for metabolic activity.□ The rigorous validation and development of *rpoB* approach provide a robust, routine and practical approach. By providing a practical means of linking community composition to physiological state, this work proposes *rpoB*Las a foundational tool for activity-resolved microbial ecology. This provides the starting point to systematic investigation of which microbial taxa drive biogeochemical processes, how active community structure responds to environmental perturbation, and whether process rates can be predicted from transcriptional profiles, questions that have remained intractable with existing marker gene approaches.

## Acknowledgement

This work is supported by a Royal□Academy of□Engineering-Scottish Water□Research□Chair (RCSRF171864) and Engineering and Physical Sciences Research Council for the Decentralised Water Technologies project (EPSCR/V030515/1).

## Data availability

Illumina NextSeq sequencing data has been uploaded to NCBI SRA under accession number PRJNA1443276 and the associated meta-data is available in the Supplementary Data 2 Sequencing_Meta_Data file. Additional nucleotide sequences of de-novo and existing primers for marker genes are available in the Supplementary Tables 3 and Supplementary data 1 Primers_Sequences file. Nucleotide and taxonomy files for the curated *rpoB* Taxonomy reference database are available in Supplementary data 3 rpoB_reference_databases (https://doi.org/10.6084/m9.figshare.33109250)

