## Supplementary Information for "Activity-resolved microbial community profiling using *rpoB* gene and transcript sequencing"

**1. Supplementary Material and methods**

**1.1 *rpoB* nucleotide sequence retrieval and databases construction:**

To retrieve *rpoB* nucleotide sequences from the NCBI datasets, a list of all available genome names was downloaded, and unique species names were compiled. One annotated genome was downloaded per unique species, prioritising complete genomes when more than one genome was available. To retrieve *rpoB* nucleotide sequences within annotated genomes, a custom R script was used, first searching for ‘gene=rpoB’ in the sequence description. When no hit was returned, alternative search criteria were used, such as ‘DNA-directed RNA polymerase subunit Beta’ or ‘RNA polymerase subunit Beta’. When the sequence for a unique species was lost during the database curation step, another genome was downloaded and used instead. *rpoB* nucleotide sequences from the GTDB database were extracted using the TIGR ID of the *rpoB* gene (TIGR02013). For the other databases, every sequence that remained after applying the search restrictions (Supplementary Table 2) were downloaded. Fasta files from individual databases were then uploaded and combined in R and duplicated sequences were removed using the *unique* function from the Biostrings package. The combined *rpoB* fasta file containing unique nucleotide sequences was then initially separated between full-length and partial-length sequences. Full-length sequences were identified using a custom R script searching for *i)* the presence of a START codon ('ATG', 'GTG', 'TTG', 'CTG', 'ATT' or 'ATC') at the beginning of the nucleotide sequence *ii)* the presence of a STOP codon (* character) at the end of the translated amino acid sequence as determined by the *translate* function from the *seqinr* package (Charif & Lobry, 2007) and *iii)* the absence of stop codon and ambiguous amino acid (letter X) within the deduced amino-acid sequence. This search for full-length sequences was done for both the sense and antisense nucleotide sequences (antisense nucleotide sequences deduced using the *reverseComplement* function form the *Biostrings* package) (Supplementary data 3). Partial-length *rpoB* nucleotide were further checked using the custom *ORF.finder* function in R (<https://github.com/Fchlt/ARC>) to retain sequences that were at least 2000bp and did not contain STOP condons in the middle of the sequence or ambiguous amino acids in their deduced amino-acid sequence. These partial-length ORF sequences were combined with the full-length rpoB fasta file to create the *rpoB* Taxonomy reference database. A custom R script was then used to find the species name (in the format ‘*genus species’, e.g. ‘Nitrosomonas aestuarii’*) for each sequence within the Taxonomy reference database. The full taxonomy for each sequence was then retrieved using the *taxonomy* function from the *myTAI* package (*db=’ncbi’, output=’classification’*), using the species name as query. If no results were found, search using a modified species name (with ‘uncultured’, ‘unknown’, ‘sp.’ removed) was used as a new query. If this new query failed too, taxonomic information was retrieved using the genus name. For a number of species, names had to be modified to reflect current taxonomic names (Supplementary Table 3). Sequences belonging to the kingdoms "Archaea","Eukaryota" and "Unknown" were removed. Once the taxonomic information was found for each unique *rpoB* sequence, a custom R script was used to retain only one representative nucleotide sequence per bacterial species. This was done to avoid biasing primers toward species containing multiple representative sequences in the combined database. This *rpoB* Primer database was further reduced to only retain sequences for which full taxonomic information was available down to species level and used for *de-novo* *rpoB* primer design/scoring and scoring (see 1.3) of existing *rpoB* primers from the literature (Supplementary Table 4).

**1.2 Reference database for other protein-coding marker genes:**

A total of 79 single copy protein-coding genes (included *rpoB*) were tested for de-novo primer design. They included all the genes listed in the UBCG2 database (kim et al 2021), for which a corresponding nucleotide fasta file was available in the GTDB database (<https://data.gtdb.aau.ecogenomic.org/releases/latest/genomic_files_all/>). To these, 19 genes involved in central cell function were added (*ATPsyn_F1gamma, atpD dnaG, dnaK, dnaN, dnaX, frr, ftsY, gyrA, hisS, holA, infC, nusG, pheT, ruvB, serS, smpB, trmD, tsf*). For all gene selected, a nucleotide fasta file was extracted from the GTDB database and curated to retain only unique full-length sequences for which full taxonomy down to species level was available following the same approach as in 1.1. Individual single copy protein-coding genes databases were then used for de-novo primer design/scoring and, when available, to score existing primers.

**1.3 Primer design and scoring:**

For *rpoB* and other protein-coding marker genes, *denovo* primer design was carried out using Primer Prospector and the *rpoB* Primer database or curated GTDB databases, respectively. First, nucleotide sequences were aligned using Mafft (Katoh et al., 2009) and primers were designed using the *generate_denovo* function from PrimerProspector, using the default parameter (length=20mers) except for the coverage which was increased to 95% of the sequences (-p 0.95) and the minimal length of the conserved 5’ region to search which was reduced to 3 nucleotides (-x 3). *De-novo* primers were initially curated considering only primers with degeneracies >384 and pairs formed based on their binding positions. For de-novo and existing primer pairs, only those for which the expected product size was compatible for both Q-PCR and Illumina sequencing (between 200 bp and 600bp) were considered.

Primers were then scored against their respective databases using the *analyse_primer.py* function from PrimerProspector. For *rpoB*, the best performing primer pairs (both de-novo and existing) were determined using in-silico PCR tests against the *rpoB* Primer database. The in-silico PCR was used to avoid using a single WS value as a threshold to consider a sequence as amplified (Okownko et al 2023). Briefly, ‘in-silico PCR’ were done by gradual increment (0.1) of the WS threshold from 0 to 5 and a sequence was considered amplified if the WS of both the forward and reverse primers was lower or equal to the threshold, considering only the sequences for which the forward primer attached before the reverse primer. For the primers designed by Bivand et al 2024, where two forward primers are used simultaneously, a sequence was considered if the WS of the reverse primer and either of the two forwards was lower than the threshold.

The three best performing primer pairs were then scored against the *rpoB* GTDB database only to compare their coverage with that of de-novo and existing primer pairs targeting other protein coding marker genes which were only scored against their respective GTDB databases. Subsequently, primer pairs were evaluated by in-silico gel electrophoresis which determined the expected amplicon size based on the matching position determined by the *analyse_primer.py* function. To do so, the binding position of the forward primer (5’ end) was subtracted to the binding position of the reverse primer (3’ end) for each sequence, again, considering only the sequences for which the forward primer attached before the reverse primer (Supplementary Figure 1).

**1.4 End-point PCR and (RT)-qPCR tests**

Selected primer pairs from the in-silico comparison were then tested by end-point PCR with a gradient of annealing temperature using soil DNA and cDNA as template. These were: the Bivand et al 2024 *rpoB* primer pair, the 1528F/2041R de-novo *rpoB* primer pair, the modified Ogier et al 201 (plus original version for comparison) primer pairs and the Barret et al 2015 *gyrB* primer pair. Based on this initial comparison, only the 1528F/2041R de-novo *rpoB* primer pair was used for Q-PCR optimisation (see main manuscript).

Q-PCR efficiency was determined using a standard curve made by 1:5 serial dilution from 10^7^ to 128 copies/µl of a synthesised DNA fragment. cDNA Q-PCR efficiency was determined by creating a 1:10 serial dilution from 10^8^ to 10^1^ of an in-vitro transcribed *rpoB* RNA fragment followed by individually reverse transcribing RNA dilutions before Q-PCR (Cholet et al., 2022 and Smith et al., 2006).

**1.5 (RT)-q-PCR and (RT)-amplicon sequencing**

DNA and RNA co-extraction for environmental (soil and sediment) samples was performed using the AllPrep PowerFecal Pro DNA/RNA Kit (Qiagen) and the ZymoBIOMICS DNA/RNA miniprep kit (Zymo Research) for BAC. RNA extractions were DNase-treated using the ezDNase kit (Thermo Fisher Scientific, UK) and the absence of DNA confirmed by negative PCR amplification of the *16S rRNA* gene after 35 cycles of PCR (Smith et al., 2006). DNA-free RNA was reverse transcribed to cDNA using the SuperScript IV kit (Thermo Fisher Scientific, UK), using random hexamer (50 ng/µl). RNA quality was evaluated using the Ramp method targeting *glnA* transcripts from the cDNA preparations (Cholet et al., 2019). *rpoB* genes and transcripts were then quantified from 1:20 diluted DNA and 1:10 diluted cDNA, respectively using the Quantitech SYBR kit (Qiagen, UK) on the ABI thermocycler (QuantStudio 3, Thermo Fisher Scientific, UK). Standard curves were constructed as described above (section 2.1.3).

Amplicon libraries were prepared following a two-step PCR amplification of the *16S rRNA* and *rpoB* gene and transcripts from extracted DNA and cDNA following Cholet et al., 2019 using the PCR primers and conditions in Table 1 and sequenced using Illumina NextSeq at (1000/2000 P1 XLEAP-SBS™ (600 Cycles) flow cell; 0.2 million reads per library) (Glasgow Polyomics <https://infraportal.org.uk/infrastructure/glasgow-polyomics>). For *16S rRNA*, ASVs were constructed using paired-end reads and the DADA2 pipeline (Callahan et al., 2016) with the *qiime dada2 denoise-paired* function. Forward and reverse primers were truncated using the *--p-trim-left-f 19* and *--p-trim-left-r 20* options, respectively. Forward and reverse reads were truncated using the *--p-trunc-len-f 270* and *--p-trunc-len-r 250* options, respectively. For *rpoB*, ASVs were constructed using DADA2 with either both forward and reverse reads (paired-end reads; *qiime dada2 denoise-paired*), testing different truncation parameters for the *--p-trunc-len-f* and *--p-trunc-len-r* options or the forward reads only (*qiime dada2 denoise-single*). The method resulting in the highest number of reads retained per samples (*qiime dada2 denoise-single*) was used for further analyses (Supplementary Table 4). For *rpoB*, only the ASVs that translated to a correct protein were retained as done in Cholet et al., (2022) (R scripts available at [https://github.com/Fchlt/ARC](https://github.com/Fchlt/ARC" \t "_blank)). Absolute abundances were inferred by multiplying relative abundances with gene or transcript numbers. ASVs taxonomy was assigned using the MIMt database (Cabezas et al., 2024) and the *rpoB* Taxonomy reference database (this study) for *16S rRNA* and *rpoB* nucleotide sequences, respectively, using the BLCA method (Gao et al., 2017). Illumina NextSeq sequencing data has been uploaded to NCBI SRA under accession number PRJNA1443276 and the associated meta-data is available in the Supplementary Data 2 Sequencing_Meta_Data file. Additional nucleotide sequences of de-novo and existing primers for marker genes are available in the Supplementary Tables 3 and Supplementary data 1 Primers_Sequences file. Nucleotide and taxonomy files for the curated *rpoB* Taxonomy reference database are available in Supplementary data 3 rpoB_reference_databases folder.

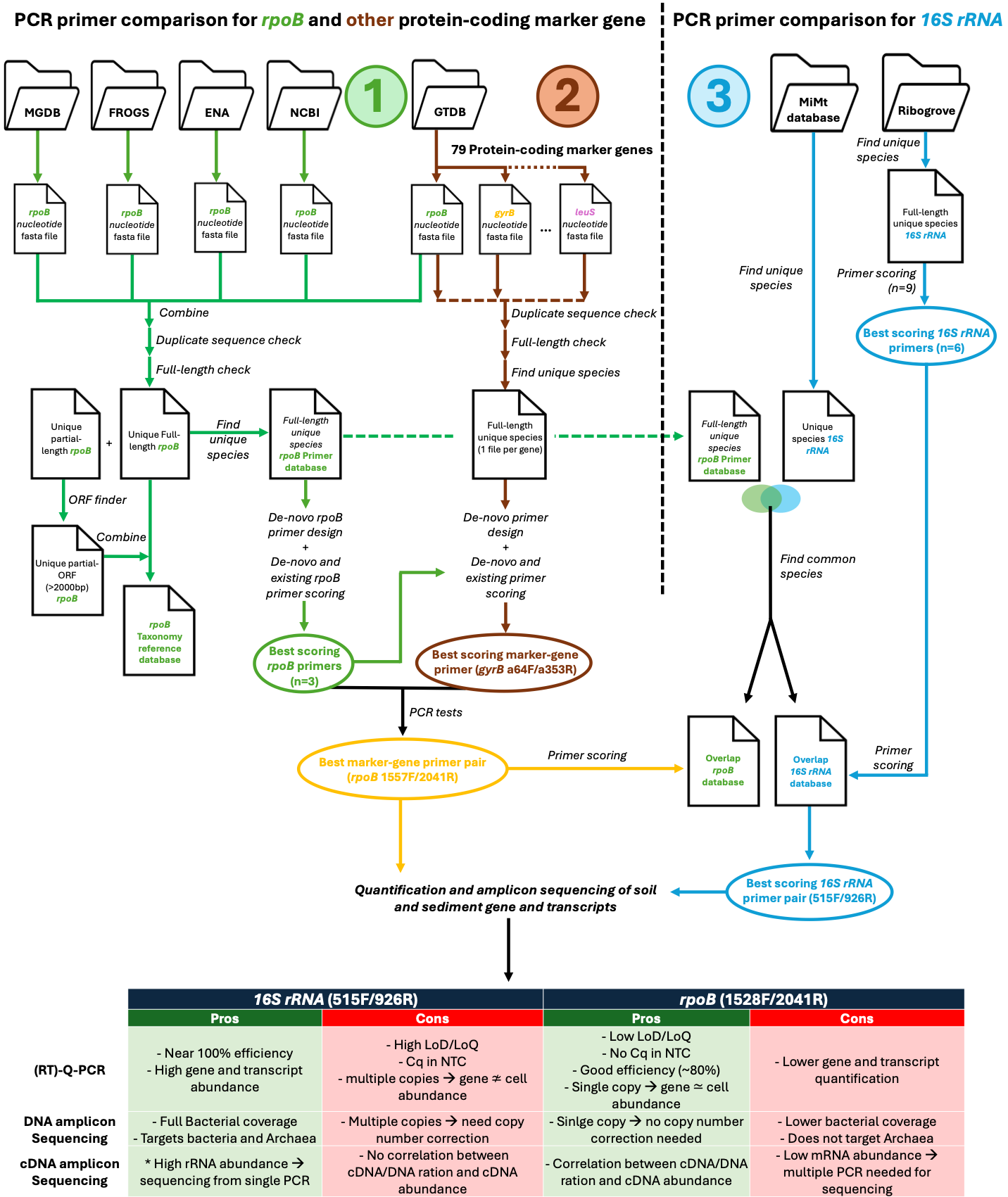

**Supplementary Figure 1. Experimental workflow and main results of this study.** The experimental workflow is divided in three main sections. The first section (green arrows) describes the process to generate the *rpoB* Taxonomy reference and the *rpoB* Primer database. The latter one is then used for de-novo *rpoB* primer design and for in-silico comparison of de-novo and existing *rpoB* primer pairs. The output of this workflow is the identification of three pairs of *rpoB* primers with best in-silico PCR performance (green oval). The second section (brown arrows) describes the process to design de-novo primers and test de-novo and existing primer pairs targeting other protein-coding marker genes and to compare their performance with the best performing rpoB primer pairs identified by workflow 1. The output of workflow 2 is the identification of one *gyrB* primer pair with better in-silico PCR performance compared to *rpoB* primer pairs (brown oval). Combined outputs from workflows 1 and 2 are used to select primers for PCR comparison, which results in the selection of a single primer pair, the *rpoB* 1528F/2041R (yellow oval). The last workflow (blue arrows) describes the process to compare the coverage of commonly used *16S rRNA* primers with that of the 1528F/2041 *rpoB* pair by creating overlap *rpoB* and *16S rRNA* databases containing common bacterial species. The output from this workflow id the identification of the *16S rRNA* 515F/926R primer pair as the best performing one. Finally, the best performing *rpoB* and *16S rRNA* are used for quantification and amplicon sequencing of total and transcriptionally active bacterial communities in soil and sediments, and the main pros and cons of both approaches are summarised.

**Supplementary Results**

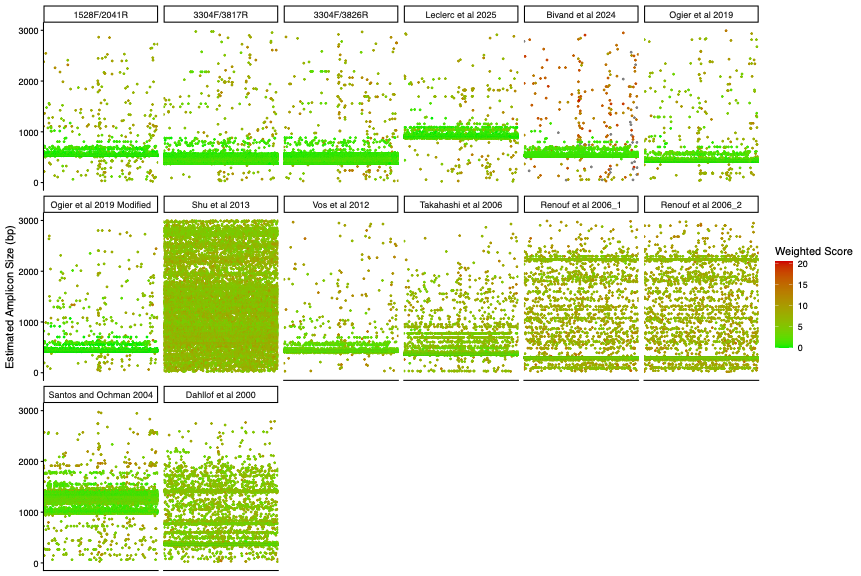

**Supplementary Figure 2. In-silico gels showing predicted amplicon size for different primer pairs targeting the *rpoB* gene.** For each primer pair, the y-axis indicates the estimated amplicon size for each sequence (individual points) and the colour of the points indicates the average WS of the forward and reverse primers.

S1.**A** *recA*

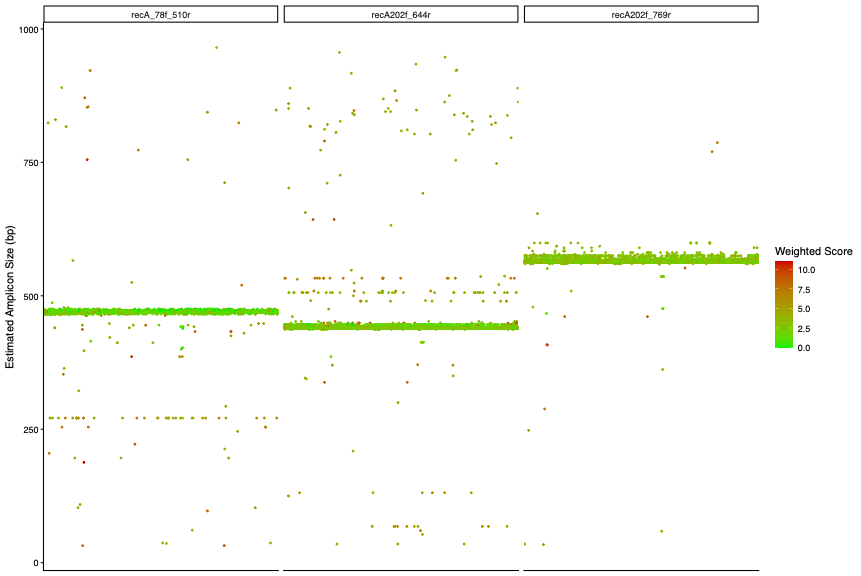

S1.B *atpD*

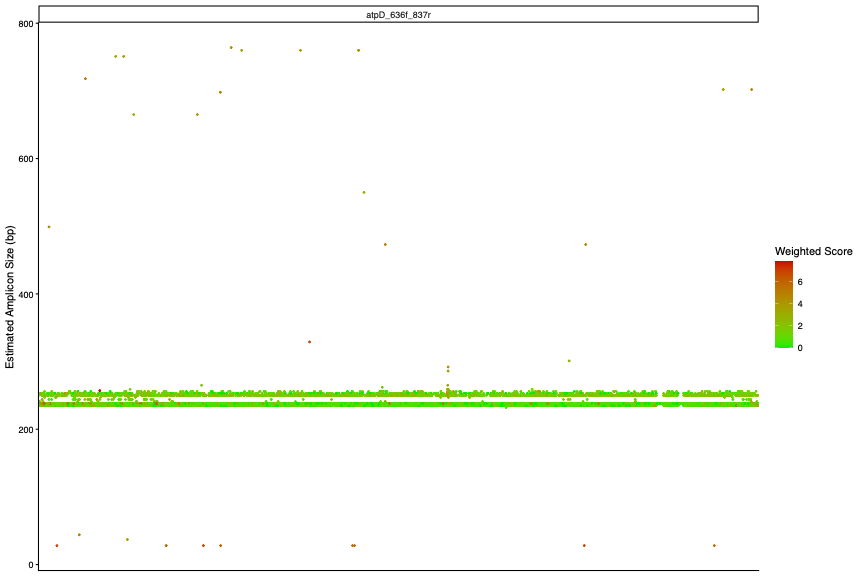

S1.C *elF2*

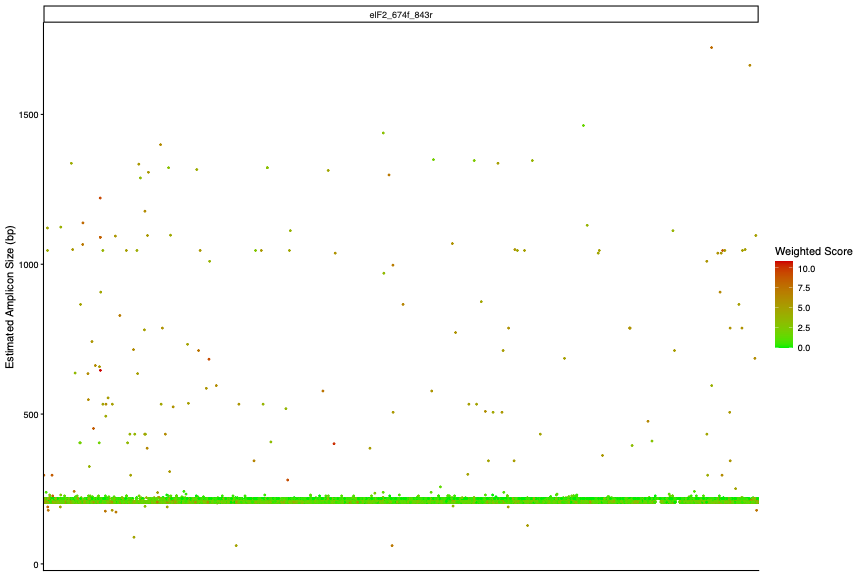

S1.D *gyrA*

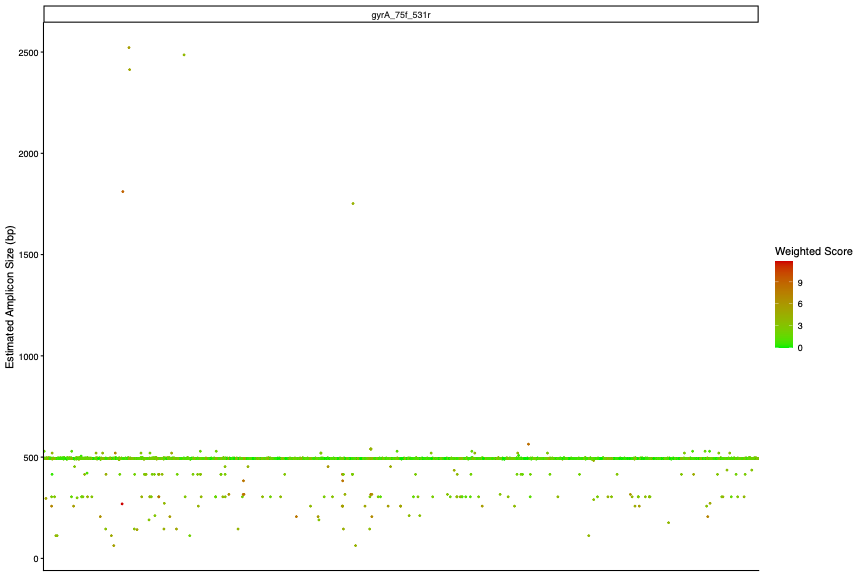

S1.E *gyrB*

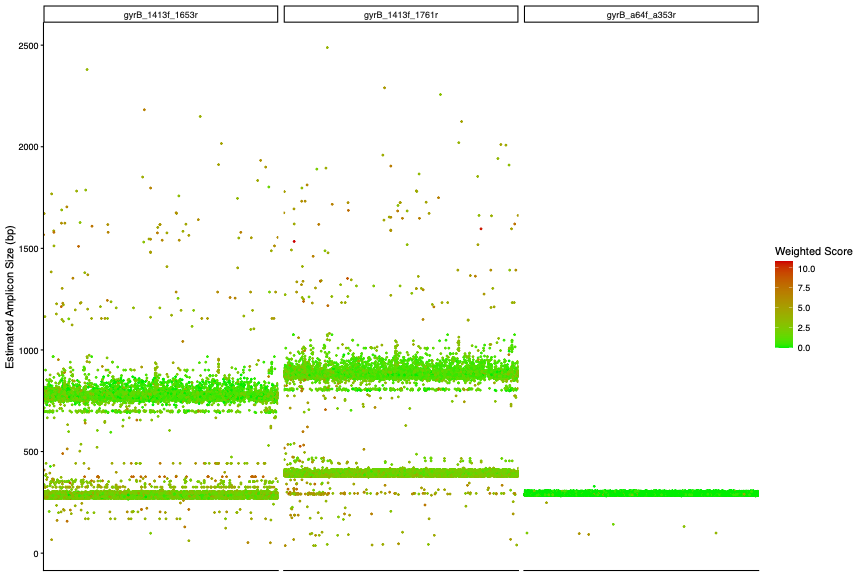

S1.F *ileS*

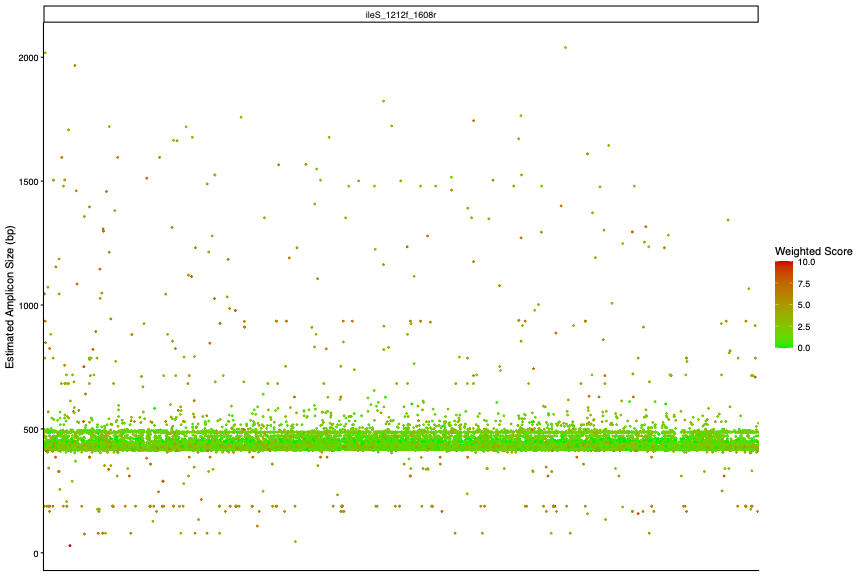

S1.G *leuS*

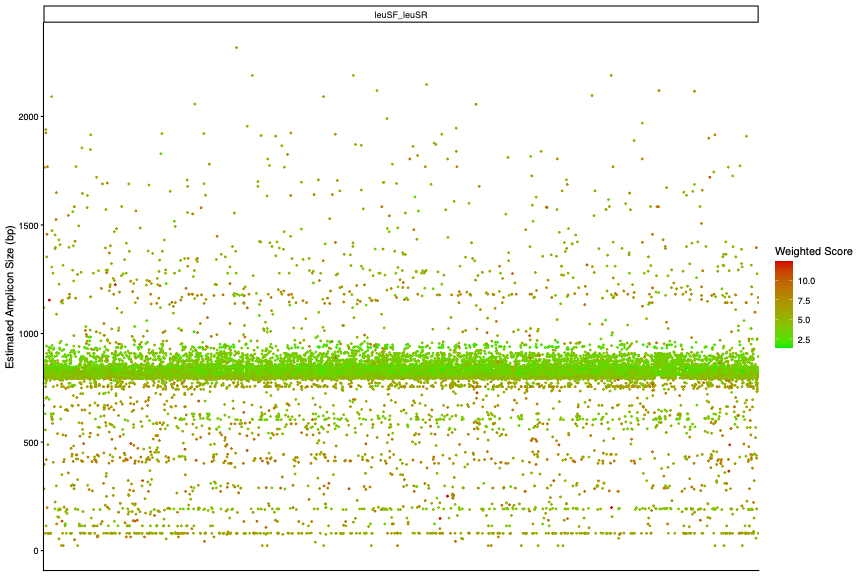

S1.H *pyrG*

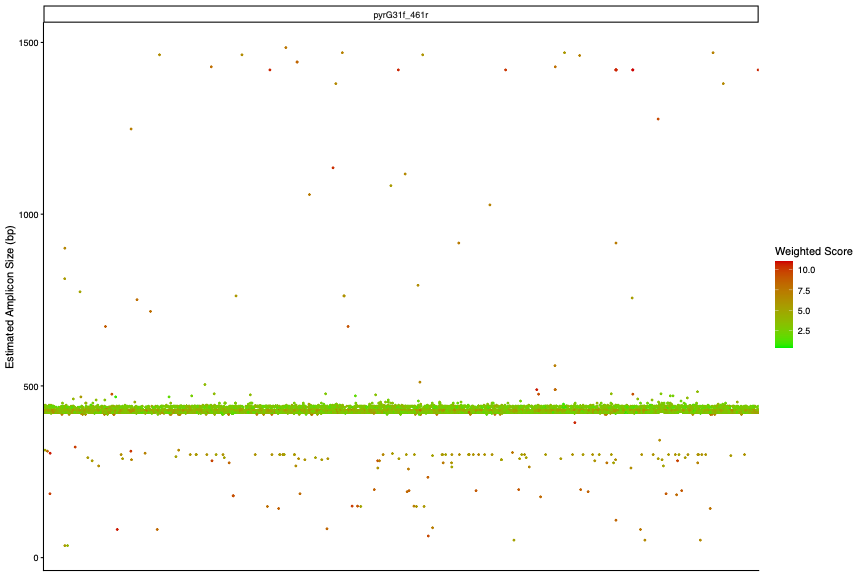

S1.I *recG*

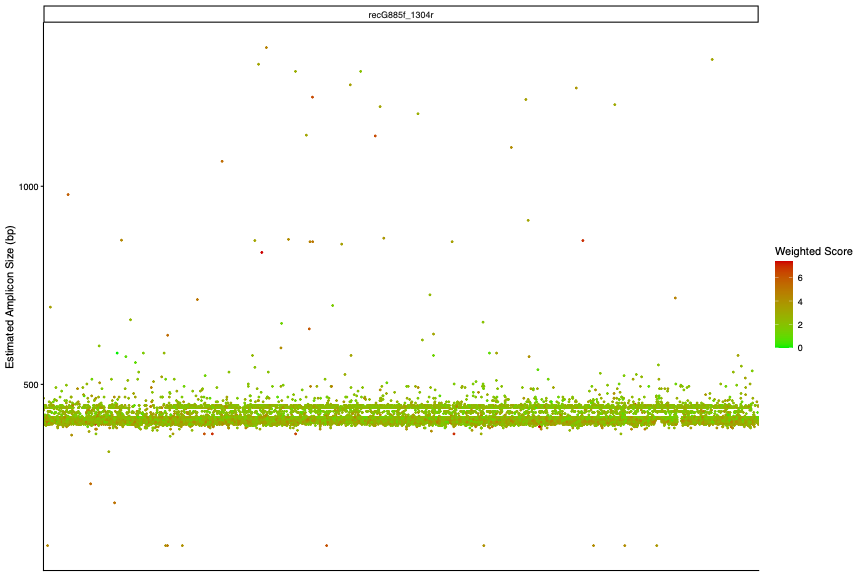

S1.J *rplB*

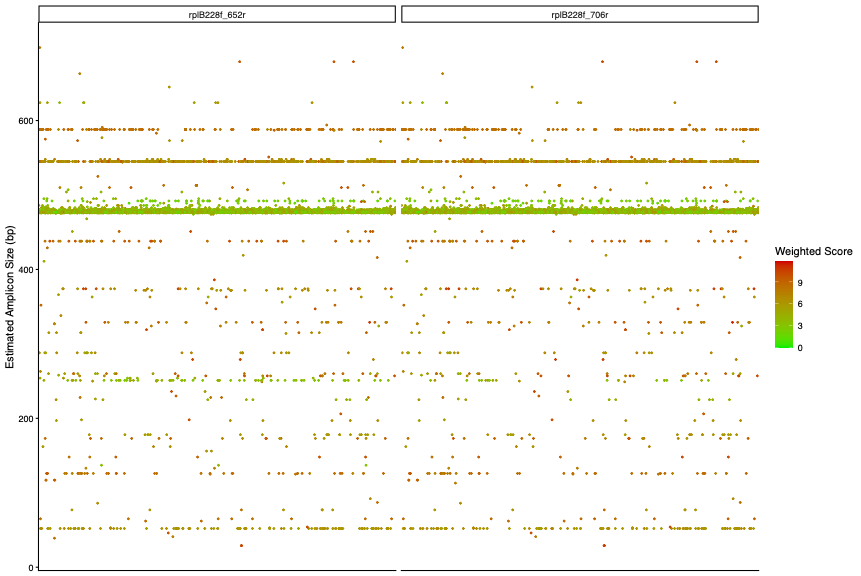

S1.K *rps2*

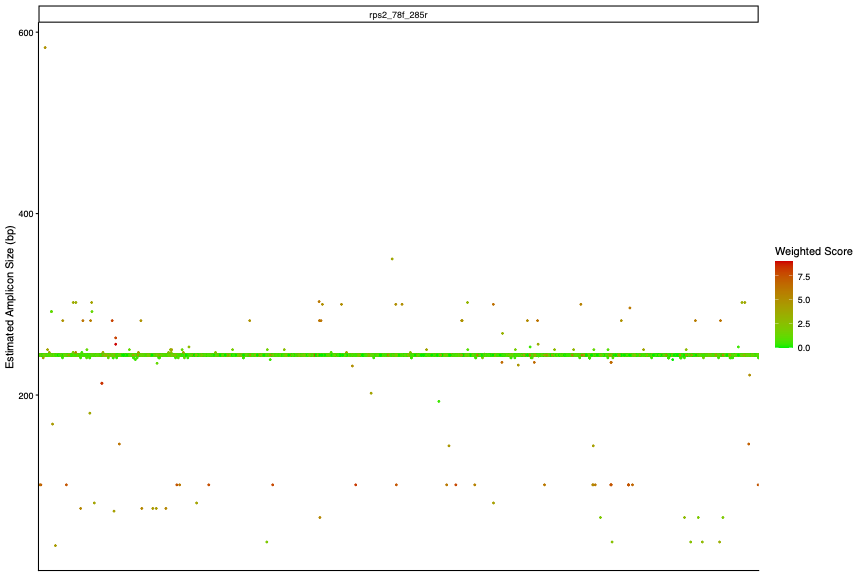

**Supplementary Figure 3. (A to K) In-silico predicted amplicon size for 11 different marker genes.** For each gene, the y-axis indicates the estimated amplicon size for each sequence (individual points). The colour of the points indicates the average WS of the forward and reverse primers.

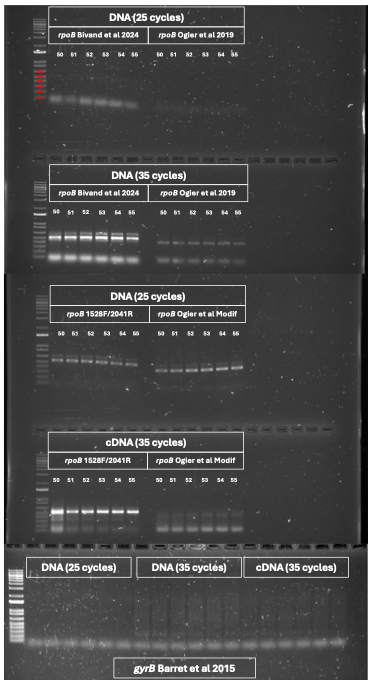

**Supplementary Figure 4.** PCR and RT-PCR amplification of best performing in silico primers for *rpoB* and *gyrB* genes

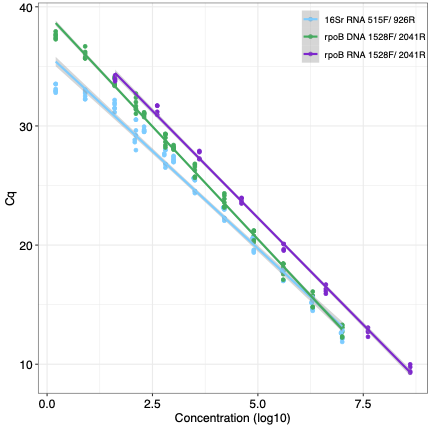

**Supplementary Figure 5.** **Q-PCR standard curves.** Standard curves for *rpoB* 1528F/2041R against standard DNA (green) and standard cDNA (purple) dilutions and for 16S rRNA 515F/926R against DNA standard dilutions (blue).

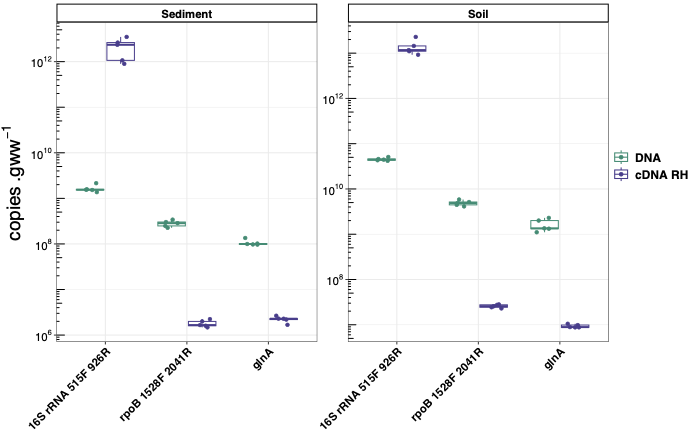

**Supplementary Figure 6.** Quantification of *16S rRNA*, *rpoB* and *glnA* genes and transcripts in sediment (left) and soil (right).

**Supplementary Table 1. List of databases used to recover *rpoB* nucleotide sequences**

| **Reference Database** | **Link** | **Search term** | **Search restriction** | **Number of Sequence** | **Number of full-ORF sequences** | **Number of partial-ORF sequences (>2,000bp)** |
| --- | --- | --- | --- | --- | --- | --- |
| **NCBI (nucleotide)** | https://[www.ncbi.nlm.nih.gov/nuccore](http://www.ncbi.nlm.nih.gov/nuccore) | *‘rpoB’* | 2000-5000bp | 7,778 | 1,508 | 1,672 |
| **ENA** | <https://www.ebi.ac.uk/ena/browser/home> | *‘RNA polymerase subunit B’* | Coding (Standard) | 9,431 | 1,986 | 486 |
| **MGDB** | <https://mbgd.nibb.ac.jp/> | *‘rpoB’* | Cluster ID 685, Bacteria | 1,930 | 1,695 | 5 |
| **FROGS** | <https://web-genobioinfo.toulouse.inrae.fr/frogs_databanks/assignation/rpoB/> | NA | NA | 362,332 | 104,028 | 4,339 |
| **NCBI datasets Bacterial genomes** | https://www.ncbi.nlm.nih.gov/datasets/genome/?taxon  =2&annotated_only=true&refseq_annotation=true&assembly_level=3:3 | Custom R script | | 193,492 | 170,550 | 4,944 |
| **GTDB** | <https://gtdb.ecogenomic.org> | TIGR02013.1 | | 638,227 | 593,651 | 40,665 |

**Supplementary Table 2. Summary of protein-coding marker genes tested in this study.**

| **Gene** | **HMM.profile** | **Function** | **Found in GTDB** | **De-novo primers found** | **Primer previously published** | **Primers matching criteria** |
| --- | --- | --- | --- | --- | --- | --- |
| alaS | TIGR00344 | Alanine-tRNA ligase | Yes |  |  |  |
| argS | TIGR00456 | Arginine-tRNA ligase | Yes |  |  |  |
| aspS | TIGR00459 | Aspartate-tRNA ligase | Yes |  |  |  |
| cgtA | TIGR02729 | GTPase ObgE/CgtA | Yes |  |  |  |
| coaE | TIGR00152 | Dephospho-CoA kinase |  |  |  |  |
| cysS | TIGR00435 | Cysteine-tRNA ligase | Yes |  |  | Yes |
| dnaA | TIGR00362 | Chromosomal replication initiator protein DnaA | Yes |  |  |  |
| dnaG | TIGR01391 | DNA primase | Yes |  |  |  |
| dnaX | TIGR02397 | DNA polymerase III subunit gamma | Yes | Yes |  |  |
| engA | TIGR03594 | GTPase Der | Yes |  |  |  |
| ffh | TIGR00959 | Signal recognition particle protein | Yes | Yes |  |  |
| fmt | TIGR00460 | Methionyl-tRNA formyltransferase | Yes |  |  |  |
| frr | TIGR00496 | Ribosome-recycling factor | Yes | Yes |  |  |
| ftsY | TIGR00064 | Signal recognition particle receptor FtsY | Yes | Yes |  |  |
| gmk | TIGR03263 | Guanylate kinase | Yes |  |  |  |
| hisS | TIGR00442 | Histidine-tRNA ligase | Yes |  |  |  |
| ileS | TIGR00392 | Isoleucine-tRNA ligase 1 | Yes | Yes | Yes | Yes |
| infB | TIGR00487 | Translation initiation factor IF-2 | Yes | Yes |  | Yes |
| infC | TIGR00168 | Translation initiation factor IF-3 | Yes |  |  |  |
| ksgA | TIGR00755 | Ribosomal RNA small subunit methyltransferase A | Yes |  |  |  |
| lepA | TIGR01393 | Elongation factor 4 | Yes | Yes | Yes |  |
| leuS | TIGR00396 | Leucine-tRNA ligase | Yes | Yes | Yes | Yes |
| ligA | TIGR00575 | DNA ligase |  |  |  |  |
| nusA | TIGR01953 | Transcription termination/antitermination protein NusA | Yes |  |  |  |
| nusG | TIGR00922 | Transcription termination/antitermination protein NusG | Yes |  |  |  |
| pgk | PF00162 | Phosphoglycerate kinase |  |  |  |  |
| pheS | TIGR00468 | Phenylalanine-tRNA ligase alpha subunit | Yes | Yes |  |  |
| pheT | TIGR00472 | Phenylalanine-tRNA ligase beta subunit | Yes | Yes |  |  |
| prfA | TIGR00019 | Peptide chain release factor 1 | Yes | Yes |  |  |
| pyrG | TIGR00337 | CTP synthase | Yes | Yes | Yes | Yes |
| recA | TIGR02012 | DNA recombination and repair protein | Yes | Yes | Yes | Yes |
| rbfA | TIGR00082 | 30S ribosome-binding factor | Yes |  |  |  |
| rnc | TIGR02191 | Ribonuclease 3 | Yes |  |  |  |
| rplA | TIGR01169 | 50S ribosomal protein L1 | Yes | Yes |  |  |
| rplB | TIGR01171 | 50S ribosomal protein L2 | Yes | Yes | Yes | Yes |
| rplC | TIGR03625 | 50S ribosomal protein L3 | Yes | Yes |  |  |
| rplD | TIGR03953 | 50S ribosomal protein L4 | Yes |  |  |  |
| rplE | PF00281 | 50S ribosomal protein L5 |  |  |  |  |
| rplF | TIGR03654 | 50S ribosomal protein L6 | Yes |  |  |  |
| rplI | TIGR00158 | 50S ribosomal protein L9 | Yes |  |  |  |
| rplJ | PF00466 | 50S ribosomal protein L10 |  |  |  |  |
| rplK | TIGR01632 | 50S ribosomal protein L11 | Yes |  |  |  |
| rplL | TIGR00855 | 50S ribosomal protein L7/L12 |  |  |  |  |
| rplM | TIGR01066 | 50S ribosomal protein L13 | Yes |  |  |  |
| rplN | TIGR01067 | 50S ribosomal protein L14 |  |  |  |  |
| rplO | TIGR01071 | 50S ribosomal protein L15 | Yes |  |  |  |
| rplP | TIGR01164 | 50S ribosomal protein L16 | Yes |  |  |  |
| rplQ | TIGR00059 | 50S ribosomal protein L17 | Yes |  |  |  |
| rplR | TIGR00060 | 50S ribosomal protein L18 |  |  |  |  |
| rplS | TIGR01024 | 50S ribosomal protein L19 |  |  |  |  |
| rplT | TIGR01032 | 50S ribosomal protein L20 | Yes |  |  |  |
| rplU | TIGR00061 | 50S ribosomal protein L21 | Yes |  |  |  |
| rplV | TIGR01044 | 50S ribosomal protein L22 | Yes |  |  |  |
| rplW | PF00276 | 50S ribosomal protein L23 |  |  |  |  |
| rplX | TIGR01079 | 50S ribosomal protein L24 | Yes |  |  |  |
| rpmA | TIGR00062 | 50S ribosomal protein L27 |  |  |  |  |
| rpmC | TIGR00012 | 50S ribosomal protein L29 |  |  |  |  |
| rpmI | TIGR00001 | 50S ribosomal protein L35 |  |  |  |  |
| rpoA | TIGR02027 | DNA-directed RNA polymerase subunit alpha | Yes |  |  |  |
| rpoB | TIGR02013 | DNA-directed RNA polymerase subunit beta | Yes | Yes | Yes | Yes |
| rpoC | TIGR02386 | DNA-directed RNA polymerase subunit beta’ | Yes | Yes |  |  |
| rpsB | TIGR01011 | 30S ribosomal protein S2 | Yes |  |  | Yes |
| rpsC | TIGR01009 | 30S ribosomal protein S3 | Yes | Yes |  |  |
| rpsD | TIGR01017 | 30S ribosomal protein S4 | Yes |  |  |  |
| rpsE | TIGR01021 | 30S ribosomal protein S5 | Yes |  |  |  |
| rpsF | TIGR00166 | 30S ribosomal protein S6 | Yes |  |  |  |
| rpsG | TIGR01029 | 30S ribosomal protein S7 | Yes | Yes |  |  |
| rpsH | PF00410 | 30S ribosomal protein S8 |  |  |  |  |
| rpsI | PF00380 | 30S ribosomal protein S9 |  |  |  |  |
| rpsJ | TIGR01049 | 30S ribosomal protein S10 |  |  |  |  |
| rpsK | TIGR03632 | 30S ribosomal protein S11 | Yes |  |  |  |
| rpsL | TIGR00981 | 30S ribosomal protein S12 |  |  |  |  |
| rpsM | TIGR03631 | 30S ribosomal protein S13 |  |  |  |  |
| rpsO | TIGR00952 | 30S ribosomal protein S15 |  |  |  |  |
| rpsP | TIGR00002 | 30S ribosomal protein S16 |  |  |  |  |
| rpsQ | TIGR03635 | 30S ribosomal protein S17 |  |  |  |  |
| rpsR | TIGR00165 | 30S ribosomal protein S18 |  |  |  |  |
| rpsS | TIGR01050 | 30S ribosomal protein S19 |  |  |  |  |
| rpsT | TIGR00029 | 30S ribosomal protein S20 | Yes |  |  |  |
| secA | TIGR00963 | Protein translocase subunit SecA | Yes |  |  | Yes |
| secG | TIGR00810 | Protein-export membrane protein SecG | Yes |  |  |  |
| secY | TIGR00967 | Protein translocase subunit SecY | Yes |  |  |  |
| serS | TIGR00414 | Serine-tRNA ligase | Yes |  |  |  |
| smpB | TIGR00086 | SsrA-binding protein | Yes |  |  |  |
| tig | TIGR00115 | Trigger factor | Yes |  |  |  |
| tilS | TIGR02432 | tRNA(Ile)-lysidine synthase | Yes |  |  |  |
| truB | TIGR00431 | tRNA pseudouridine synthase B | Yes |  |  |  |
| tsaD | TIGR03723 | tRNA N6-adenosine threonylcarbamoyltransferase | Yes |  |  |  |
| tsf | TIGR00116 | Elongation factor Ts | Yes | Yes |  |  |
| uvrB | TIGR00631 | UvrABC system protein B | Yes |  |  | Yes |
| ybeY | TIGR00043 | Endoribonuclease YbeY | Yes |  |  |  |
| ychF | TIGR00092 | Ribosome-binding ATPase YchF | Yes |  |  |  |
| Der | TIGR03594 | ribosome biogenesis GTPase Der | Yes | Yes |  |  |
| atpD | TIGR01039 | ATP synthase F1, beta subunit | Yes | Yes |  | Yes |
| atp_F1gamma | TIGR01146 | ATPsyn_F1gamma | Yes | Yes |  |  |
| dnaN | TIGR00663 | DNA polymerase III, beta subunit | Yes |  |  |  |
| dnaK | TIGR02350 | chaperone protein DnaK | Yes | Yes |  |  |
| gyrA | TIGR01063 | DNA gyrase, A subunit | Yes | Yes |  | Yes |
| gyrB | TIGR01059 | DNA gyrase, B subunit | Yes | Yes | Yes | Yes |
| recG | TIGR00643 | ATP-dependent DNA helicase RecG | Yes |  | Yes | Yes |
| ruvB | TIGR00635 | Holliday junction DNA helicase RuvB | Yes | Yes |  |  |
| trmD | TIGR00088 | tRNA (guanine(37)-N(1))-methyltransferase | Yes | Yes |  |  |
| holA | TIGR01128 | DNA polymerase III, delta subunit | Yes |  |  |  |

**Supplementary Table 3. List of *rpoB*, *16S rRNA* and other protein-coding marker gene primers for in-silico and PCR comparison.** Start and end positions were determined against respective *E.coli* genes. Degeneracy numbers correspond to the product of all degeneracies present in the primer sequence (N=4, B/D/H/V=3, Y/W/S/M/K/R=2). Primer pairs tested in PCR are highlighted with the same colour. Tests: IS= *in-silico* PCR; PCR= end-point (RT)-PCR. Primers in red were not tested because they were expected to produce amplicon too long for Q-PCR and Illumina amplicon sequencing.

| **Primer** | **Target gene** | **Orientation** | **Sequence (5' à 3')** | **Degeneracies** | **Start** | **End** | **Tests** | **Reference** |
| --- | --- | --- | --- | --- | --- | --- | --- | --- |
| 1528f | Bacterial RNA polymerase subunit B (*rpoB*) | Forward | CAGYTGTCBCAGTTYATGGA | 12 | 1528 | 1547 | IS/PCR | This Study (de-novo) |
| 3304f |  |  | GGYGTRCCKTSBCGKATGAA | 96 | 3304 | 3323 | IS |  |
| 2041r |  | Reverse | CGYTGCATGTTSGMDCCCAT | 24 | 2041 | 2060 | IS/PCR |  |
| 3817r |  |  | TCMAGYGCCCAVACYTCCAT | 24 | 3817 | 3836 | IS |  |
| 3826r |  |  | CCRTADGCYTCMAGYGCCCA | 48 | 3826 | 3845 |  |  |
| rpoB1f |  | Forward | CGTGCACCCCACCCAYTAYGGNMG | 32 | 1647 | 1670 |  | Vos et al 2012 |
| rpoB1r |  | Reverse | CACGGCCTGCCKYTGCATRTT | 8 | 2050 | 2070 |  |  |
| rpoB4f |  | Forward | CGAACATCGGTCTGATCAACTC | 0 | 1700 | 1723 |  | Takahashi et al 2006 |
| rpoB2r |  | Reverse | GTTGCATGTTCGCACCCAT | 0 | 2041 | 2059 |  |  |
| rpoB1698f |  | Forward | AACATCGGTTTGATCAAC | 0 | 1702 | 1719 |  | Dahllöf et al 2000 |
| rpoB2041r |  | Reverse | CGTTGCATGTTGGTACCCAT | 0 | 2041 | 2060 |  |  |
| Univ_rpoB_F_deg |  | Forward | GGYTWYGAAGTNCGHGACGTDCA | 288 | 1630 | 1653 | IS/PCR | Ogier et al 2019 |
| Univ_rpoB_R_deg |  | Reverse | TGACGYTGCATGTTBGMRCCCATMA | 48 | 2039 | 2063 |  |  |
| rpoB_ModF |  | Forward | GGYTTYGARGTBCGYGACGT | 48 | 1630 | 1649 | IS/PCR | This study; modified from Ogier et al 2019 |
| rpoB_ModR |  | Reverse | CGYTGCATGTTSGMDCCCAT | 24 | 2041 | 2060 |  |  |
| rpoBDP0r |  | Reverse | CNGCYTGDCKYTKCATRTTNNNNCCCAT | 98304 | 2041 | 2068 | IS/PCR | Bivand et al 2024 |
| rpoBDP01f |  | Forward | TCNCARTTYATGGAYCANNHNAAYCC | 12288 | 1534 | 1559 |  |  |
| rpoBDP02f |  |  | AGNCARTTYATGGAYCANNHNAAYCC | 12288 | 1534 | 1559 |  |  |
| rpoBoutf |  | Forward | CAGYTDTCNCARTTYATGGAYCA | 192 | 1528 | 1550 | IS | Leclerc et al 2025 |
| rpoBoutr |  | Reverse | AGTTRTARCCDTYCCANGKCAT | 192 | 2413 | 2434 |  |  |
| rpoB1 |  | Forward | ATTGACCACTTGGGTAACCGTCG | 0 | 1333 | 1355 |  | Renouf et al 2006 |
| rpoB1o |  |  | ATCGATCACTTAGGCAATCGTCG | 0 | 1333 | 1355 |  |  |
| rpoB2 |  | Reverse | ACGATCACGGGTCAAACCACC | 0 | 1606 | 1626 |  |  |
| 27f | Prokaryotic small-subunit Ribosomal RNA gene (*16S rRNA*) | Forward | AGAGTTTGATCCTGGCTCAG | 0 | 8 | 27 | IS | Mesa et al 2017 |
| 63f |  |  | CAGGCCTAACACATGCAAGTC | 0 | 43 | 63 |  |  |
| 338r |  | Reverse | TGCTGCCTCCCGTAGGAGT | 0 | 338 | 356 |  | Mesa et al 2017 |
| V2f |  | Forward | AGTGGCGGACGGGTGAGTAA | 0 | 101 | 120 |  | Will et al 2010 |
| V3r |  | Reverse | CCGCGGCTGCTGGCAC | 0 | 515 | 530 |  | Sahm et al 2013 |
| 341f |  | Forward | CCTACGGGNGGCWGCAG | 8 | 341 | 357 |  | Klindworth et al 2013 |
| 785r |  | Reverse | GACTACHVGGGTATCTAATCC | 9 | 785 | 805 |  |  |
| 806RBr |  |  | GGACTACNVGGGTWTCTAAT | 24 | 787 | 806 |  | Apprill et al 2015 |
| 515FYf |  | Forward | GTGYCAGCMGCCGCGGTAA | 4 | 515 | 533 | IS/PCR | Parada et al 2016 |
| 926r |  | Reverse | CCGYCAATTYMTTTRAGTTT | 16 | 907 | 926 |  | Quince et al 2011 |
| 1369f |  | Forward | CGGTGAATACGTTCYCGG | 2 | 1369 | 1386 | IS | Suzuki et al 2000 |
| 1492r |  | Reverse | GGWTACCTTGTTACGACTT | 2 | 1492 | 1510 |  |  |
| B969f |  | Forward | ACGCGHNRAACCTTACC | 24 | 969 | 985 |  | Comeau et al 2017 |
| BA1406r |  | Reverse | ACGGGCRGTGWGTRCAA | 8 | 1390 | 1406 |  |  |
| 78f | *rps2* | Forward | GYHRYTGGAAYCCRAARATG | 192 | 78 | 98 | IS | This Study |
| 285r |  | Reverse | GTSARCRTRCCRCCBARCCA | 192 | 285 | 305 |  |  |
| 78f | *recA* | Forward | TYGGYAARGGYKCSRTCATG | 128 | 78 | 98 | IS | This Study |
| 510r |  | Reverse | TTRCGCARSGCYTGRSWCAT | 128 | 510 | 530 |  |  |
| recABDUP1 |  | Forward | CCCGAGTCCTCCGGNAARACNAC | 32 | 202 | 225 |  | Santos and Ochman 2004 |
| recABGDN2 |  | Reverse | CGTTGCCGCCGGKNGTNRYYTC | 256 | 622 | 644 |  |  |
| recABHDN1 |  | Reverse | GAAGGGTGGGGCCANYTTRTTYTT | 32 | 745 | 769 |  |  |
| 1413f | *gyrB* | Forward | AYCAYARVATCRTYMTCATG | 192 | 1413 | 1433 | IS | This Study |
| 1653r |  | Reverse | CAMARYTSYTCSGSRTTCAT | 256 | 1653 | 1673 |  |  |
| 1761r |  | Reverse | GGYTCVACBTCRTCRCCCAT | 72 | 1761 | 1781 |  |  |
| a64f |  | Forward | MGNCCNGSNATGTAYATHGG | 1536 | 64 | 84 | IS/PCR | Barret et al 2015 |
| a353r |  | Reverse | ACNCCRTGNARDCCDCCNGA | 2304 | 334 | 354 |  |  |
| Up1f |  | Forward | CAYGCNGGNGGNAARTTYGA | 512 | 296 | 315 | IS | Yamamoto and Harayama 1995 |
| Up2r |  | Reverse | CCRTCNACRTCNGCRTCNGTCAT | 512 | 1487 | 1509 |  |  |
| gyrBBAUP2 |  | Forward | GCGGAAGCGGCCNGSNATGTA | 32 | 57 | 78 |  | Santos and Ochman 2004 |
| gyrBBNDN1 |  | Reverse | CCGTCCACGTCGGCRTCNGYCAT | 16 | 1486 | 1509 |  |  |
| 674f | *infB* | Forward | CCRGTBGTBACSRTCATGGG | 72 | 674 | 694 | IS | This Study |
| 843r |  | Reverse | TKVGCRCCRCGGGCACGCAT | 24 | 843 | 863 |  |  |
| 636f | *atpD* | Forward | CSMTSGTBTWYGGYCARATG | 384 | 636 | 656 | IS | This Study |
| 837r |  | Reverse | CGYTCYTGCARYDSRCCCAT | 192 | 837 | 857 |  |  |
| 1212f | *ileS* | Forward | ACWSCTAYCCRCAYTGCTGG | 32 | 1212 | 1232 | IS | This Study |
| 1608r |  | Reverse | RTSGMRCCSGARTCRAACCA | 128 | 1608 | 1628 |  |  |
| ileSBCUP1 |  | Forward | GCCCGGCTGGGAYWSNCAYGG | 64 | 276 | 296 |  | Santos and Ochman 2004 |
| ileSBKDN1 |  | Reverse | TGGAGCCGGAGTCAWCCANMMNTC | 128 | 1594 | 1619 |  |  |
| 75f | *gyrA* | Forward | SCTAYMTCGAYTAYGCSATG | 64 | 75 | 95 | IS | This Study |
| 531r |  | Reverse | GGCGGRATRTTGGTSGCCAT | 8 | 531 | 551 |  |  |
| leuSF | *leuS* | Forward | GAGACCGTGCTGGCCAYGARSARRT | 32 | 1273 | 1293 | IS | Santos and Ochman 2004 |
| leuSBKDN1 |  | Reverse | GGGGCAGCCCCARWANCKYT | 64 | 2476 | 2496 |  |  |
| 1111f | *urvB* | Forward | BCCGCARRTYSGCGSSATGT | 192 | 1111 | 1131 | IS | This Study |
| 1434r |  | Reverse | TCGGTSARRTCYTCSGCCAT | 32 | 1434 | 1454 |  |  |
| 1160 | *secA* | Forward | RAVAARCTSKCSGGYATGAC | 192 | 1160 | 1180 | IS |  |
| 1506 |  | Reverse | TCSGTRCCRCGRCCGGCCAT | 16 | 1506 | 1526 |  |  |
| 648f | *cysS* | Forward | AYMTYGARTGYTCSGCSATG | 128 | 648 | 668 | IS |  |
| 813r |  | Reverse | TTGCCSARSGAYTTSSWCAT | 128 | 813 | 833 |  |  |
| rplBBDUP1 | *rplB* | Forward | CAAGGTGGAGCGCATCSANTAYGAYCC | 32 | 228 | 255 | IS | Santos and Ochman 2004 |
| rplBBHDN1 |  | Forward | GCCGCCGCCGWDNGGRTGRTC | 96 | 632 | 652 |  |  |
| rplBR |  | Reverse | CGCCGCCGCCGWRNGGRTGRTC | 64 | 685 | 706 |  |  |
| recGBHUP2 | *recG* | Forward | GGGCGACGTGGGCDSNGGNAARAC | 192 | 885 | 909 | IS |  |
| recGBMDN1 |  | Reverse | GGGTCCGGGGGATNGGNGTNGC | 64 | 1282 | 1304 |  |  |
| pyrGBAUP1 | *pyrG* | Forward | GGCGTGGTGTCCTCCNTNGGNAARGG | 128 | 31 | 57 | IS |  |
| pyrGBDDN2 |  | Reverse | GGAAGGGCAGGCACTCNATRTCNCCNA | 128 | 434 | 461 |  |  |

**Supplementary Table 4.** **Determination of optimal parameters to generate *rpoB* ASVs.** For each test, the number of bases removed from the forward (X) and reverse (Y) are indicated by X_Y, respectively. The method used for comparison of sequencing results with *16S rRNA* is highlighted in green.

| **Sample Type** | **Target Nucleic Acid** | **Average % reads retained 0_40 bp** | **Average % reads retained 0_45 bp** | **Average % reads retained 0_50 bp** | **Average % reads retained 0_55 bp** | **Average % reads retained 5_45 bp** | **Average % reads retained 5_50 bp** | **Average % reads retained dada_single (no bases trimmed)** |
| --- | --- | --- | --- | --- | --- | --- | --- | --- |
| **Sediment** | **DNA** | 72.56 | 72.88 | 82.66 | 72.9 | 72.76 | 72.76 | 93.774 |
|  | **cDNA** | 64.5 | 65.12 | 70.56 | 65.5 | 65.28 | 65.48 | 88.238 |
| **Soil** | **DNA** | 82.4 | 82.66 | 82.94 | 81.54 | 82.42 | 81.4 | 94.21 |
|  | **cDNA** | 57.58 | 58.56 | 59.56 | 58.72 | 58.5 | 58.68 | 82.936 |

**Supplementary Table 5. Changes in slope for the *rpoB* and 16S rRNA assays between successive concentrations.**

| **Points** | **Slope** | **Efficiency** | **Target** |
| --- | --- | --- | --- |
| 10^7^ 🡪 2.10^6^ | -3.73 | 85.33 | *16SrRNA* 515/926 |
| 2.10^6^ 🡪 4.10^5^ | -3.43 | 95.63 |  |
| 4.10^5^🡪8.10^4^ | -3.57 | 90.72 |  |
| 8.10^4^🡪1.6.10^4^ | -3.78 | 83.92 |  |
| 1.6.10^4^🡪3.2.10^3^ | -3.36 | 98.24 |  |
| 3.2.10^3^🡪10^3^ | -4.62 | 64.61 |  |
| 10^3^🡪6.4.10^2^ | -0.08 | 1.3.10^15^ |  |
| 6.4.10^2^🡪2.10^2^ | -4.81 | 61.47 |  |
| 2.10^2^🡪128 | 0.33 | -99.90 |  |
| 128🡪40 | -4.16 | 73.83 |  |
| 40🡪8 | -1.34 | 460.48 |  |
| 8🡪1.6 | -0.75 | 2070.07 |  |
| 10^7^ 🡪 2.10^6^ | -3.69 | 86.51 | *rpoB* 1557F/ModR |
| 2.10^6^ 🡪 4.10^5^ | -3.64 | 88.10 |  |
| 4.10^5^🡪8.10^4^ | -4.00 | 77.75 |  |
| 8.10^4^🡪1.6.10^4^ | -4.08 | 75.82 |  |
| 1.6.10^4^🡪3.2.10^3^ | -3.68 | 86.94 |  |
| 3.2.10^3^🡪10^3^ | -3.90 | 80.59 |  |
| 10^3^🡪6.4.10^2^ | -3.61 | 89.09 |  |
| 6.4.10^2^🡪2.10^2^ | -4.17 | 73.61 |  |
| 2.10^2^🡪128 | -4.19 | 73.34 |  |
| 128🡪40 | -3.63 | 88.49 |  |
| 40🡪8 | -3.40 | 96.99 |  |
| 8🡪1.6 | -2.24 | 179.41 |  |

**Supplementary Table 6. Modified *rpoB* taxon’s names.**

| **Old Name** | **New Name** | **Taxonomic level** |
| --- | --- | --- |
| Izimaplasma | Izemoplasma | Genus |
| Baumannia | Candidatus Palibaumannia |  |
| Ishikawaella | Candidatus Ishikawella |  |
| Tremblaya | Candidatus Tremblayella |  |
| Ruthia | Candidatus Ruthturnera |  |
| Evansia | Candidatus Johnevansia |  |
| Brevefilum | Candidatus Brevifilum |  |
| Promineofilum | Candidatus Promineifilum |  |
| Blochmannia | Candidatus Blochmanniella |  |
| Hamiltonella | Candidatus Williamhamiltonella |  |
| Kinetoplastibacterium | Candidatus Kinetoplastidibacterium |  |
| Arthromitus | Candidatus Neoarthromitus |  |
| Candidatus Sulcia | Candidatus Karelsulcia |  |
| Sphingobacteruim | Sphingobacterium |  |
| Candidatus Vallotia | Candidatus Vallotiella |  |
| Candidatus Neptunochlamydia | Candidatus Neptunichlamydia |  |
| Candidatus Dwaynia | Candidatus Dwaynesavagella |  |
| Candidatus Endonucleobacter | Candidatus Endonucleibacter |  |
| Candidatus Protistobacter | Candidatus Protistibacter |  |
| Candidatus Micropelagos | Candidatus Micropelagius |  |
| Allitabrizicola | Alitabrizicola |  |
| Candidatus Megaira | Candidatus Megaera |  |
| Candidatus Scubalenecus | Candidatus Scybalenecus |  |
| Candidatus Microthrix | Candidatus Neomicrothrix |  |
| Candidatus Moanabacter | Candidatus Moanibacter |  |
| Candidatus Carbobacillus | Candidatus Carbonibacillus |  |
| Candidatus Lumbricidophila | Candidatus Lumbricidiphila |  |
| Candidatus Magnetoovum | Candidatus Magnetovum |  |
| Terrarubrum | Terrirubrum |  |
| Streptoverticillium | Streptomyces |  |
| Candidatus Koribacter | Candidatus Korobacter |  |
| Hassalia | Hassallia |  |
| Candidatus Galacturonibacter | Candidatus Galacturonatibacter |  |
| Alisedimentitalea | Aliisedimentitalea |  |
| Candidatus Microthrix | Candidatus Neomicrothrix |  |
| Candidatus Contendobacter | Candidatus Contendibacter |  |
| Candidatus Microthrix | Candidatus Neomicrothrix |  |
| Tetrasphaera | Nostocoides |  |
| Candidatus Endoecteinascidia | Candidatus Endecteinascidia |  |
| Candidatus Vesicomyosocius | Candidatus Vesicomyidisocius |  |
| Candidatus Thermofonsia | Candidatus Thermofontia |  |
| Candidatus Aminicenantes | Candidatus Aminicenantota |  |
| Candidatus Aerophobetes | Candidatus Aerophobota |  |
| Candidatus Coatesbacteria | Candidatus Coatesiibacteriota |  |
| Candidatus Rokubacteria | Candidatus Rokuibacteriota |  |
| Candidatus Omnitrophica | Candidatus Omnitrophota |  |
| Candidatus Phaeomarinobacter | Candidatus Phaeomarinibacter |  |
| Muricauda | Allomuricauda |  |
| Sinirhodobacter | Paenirhodobacter |  |
| Fournierella | Allofournierella |  |
| Ruthia magnifica | Candidatus Ruthturnera calyptogenae | Species |
| Blochmannia floridanus | Candidatus Blochmanniella floridana |  |
| Hamiltonella defensa | Candidatus Williamhamiltonella defendens |  |
| Kinetoplastibacterium galatii | Candidatus Kinetoplastidibacterium galati |  |
| Vesicomyosocius okutanii | Candidatus Vesicomyidisocius calyptogenae |  |
| Candidatus Protistobacter heckmanni | Candidatus Protistibacter heckmannii |  |
| Candidatus Chryseobacterium massiliae | Candidatus Chryseobacterium massiliense |  |
| Candidatus Magnetoovum chiemensis | Candidatus Magnetovum chiemense |  |
| Candidatus Contendobacter odensis | Candidatus Contendibacter odensensis |  |
| Candidatus Endoecteinascidia frumentensis | Candidatus Endecteinascidia fromenterensis |  |
| Candidatus Kinetoplastidibacterium blastocrithidii | Candidatus Kinetoplastidibacterium blastocrithidiae |  |
| Candidatus Kinetoplastidibacterium desouzaii | Candidatus Kinetoplastidibacterium desouzai |  |
| Candidatus Kinetoplastidibacterium crithidii | Candidatus Kinetoplastidibacterium crithidiae |  |
| Candidatus Kinetoplastidibacterium oncopeltii | Candidatus Kinetoplastidibacterium stringomonadis |  |
| Candidatus Kinetoplastidibacterium sorsogonicusi | Candidatus Kinetoplastidibacterium stringomonadis |  |
| Caldiarchaeum subterraneum | Candidatus Caldarchaeum subterraneum |  |
| Phaeomarinobacter ectocarpi | Candidatus Phaeomarinibacter ectocarpi |  |
| Candidatus Prevotella | Prevotella conceptionensis |  |
| Weisella paramesenteroides | Weissella paramesenteroides |  |
| Candidatus Promineifilum glycogenicum | Candidatus Promineofilum glycogenicum |  |
| Candidatus Rickettsia tasmanensis | Candidatus Rickettsia tasmaniensis |  |
| Rickettsia barbariae | Candidatus Rickettsia barbarica |  |
| Candidatus Arcanobacter lacustris | Candidatus Arcanibacter lacustris |  |
| Bactericera trigonica | Sodalis-like symbiont of Bactericera trigonica |  |
| Lysobacter terrestris | Agrilutibacter terrestris |  |
| Candidatus Rickettsia barbariae | Candidatus Rickettsia barbarica |  |
| Nitrosotalea devanaterra | Nitrosotalea devaniterrae |  |
| Lactobacillus plantarum | Lactiplantibacillus plantarum |  |
| Nitrosomarinus catalina | Candidatus Nitrosomarinus catalinensis |  |
| Tokpelaia hoelldoblerii | Candidatus Tokpelaia hoelldobleri |  |
| Propionibacterium propionicum | Arachnia propionica |  |
| Methanomethylophilus alvus | Methanomethylophilus alvi |  |
| Caedibacter acanthamoebae | Candidatus Paracaedimonas acanthamoebae |  |
| Desulfobacterium autotrophicum | Desulforapulum autotrophicum |  |
| Pseudomonas stutzeri | Stutzerimonas stutzeri |  |
| Staphylococcus sciuri | Mammaliicoccus sciuri |  |
| Sulcia muelleri | Candidatus Karelsulcia muelleri |  |
| Bacillus halodurans | Halalkalibacterium halodurans |  |
| Candidatus Neoehrlichia lotoris | Candidatus Neoehrlichia procyonis |  |
| Candidatus Photodesmus blepharus | Candidatus Photodesmus blepharonis |  |
| Hungatella xylanolytica | Lacrimispora xylanisolvens |  |
| Rhodobacter xinxiangensis | Falsirhodobacter xinxiangensis |  |
| Candidatus Sodalis pierantonius | Candidatus Sodalis pierantonii |  |
| Vreelandella utahensis | Vreelandella halophila |  |
| Thermostichus lividus | Parathermosynechococcus lividus |  |
| Candidatus Paracaedibacter acanthamoebae | Candidatus Odyssella acanthamoebae |  |
| Candidatus Photodesmus blepharus | Candidatus Photodesmus blepharonis |  |
| Hungatella xylanolytica | Lacrimispora xylanisolvens |  |
| Achromobacter aestuarii | Schauerella aestuarii |  |
| Zongyanglinia marina | Parasedimentitalea maritima |  |
| Spiribacter halobius | Sediminicurvatus halobius |  |
| Lysobacter solisilvae | Agrilutibacter solisilvae |  |
| Paenimyroides aquimaris | Paenimyroides marinum |  |
| Oscillatoria nigro-viridis | Phormidium nigroviride |  |
| Pedobacter xinjiangensis | Desertivirga xinjiangensis |  |
| Clostridium clostridioforme | Enterocloster clostridioformis |  |
| Candidatus Bandiella woodruffii | Candidatus Bandiella euplotis |  |
| Candidatus Fokinia cryptica | Candidatus Fokinia crypta |  |
| Rhodobacter kunshanensis | Paragemmobacter kunshanensis |  |
| Pseudalkalibacillus caeni | Exobacillus caeni |  |
| Candidatus Rhodobacter lobularis | Candidatus Rhodobacter oscarellae |  |
| Defluviimonas denitrificans | Albidovulum denitrificans |  |
| Candidatus Endomicrobium trichonymphae | Candidatus Endomicrobiellum trichonymphae |  |
| Pasteurella langaaensis | Alitibacter langaaensis |  |
| Rhodobaca bogoriensis | Roseinatronobacter bogoriensis |  |
| Candidatus Atelocyanobacterium thalassa | Candidatus Atelocyanobacterium thalassae |  |
| Blochmannia endosymbiont of Camponotus | unclassified Candidatus Blochmanniella |  |
| Blochmannia endosymbiont of Polyrhachis | unclassified Candidatus Blochmanniella |  |
| Rhodococcus corynebacterioides | Rhodococcoides corynebacterioides |  |
| Peptoniphilus nemausensis | Aedoeadaptatus nemausensis |  |
| Roseibaca domitiana | Roseinatronobacter domitianus |  |
| Salinibacterium sedimenticola | Homoserinimonas sedimenticola |  |
| Rhizobium rhizolycopersici | Mycoplana rhizolycopersici |  |
| Rubrivivax pictus | Pseudaquabacterium pictum |  |
| Microbacterium humi | Paramicrobacterium humi |  |
| Pseudoclavibacter alba | Pseudoclavibacter albus |  |
| Enterovibrio luxaltus | Candidatus Enterovibrio altilux |  |
| Sphingosinicella ginsenosidimutans | Allosphingosinicella ginsenosidimutans |  |
| Candidatus Epulopiscium viviparus | Candidatus Epulonipiscium viviparus |  |
| Bacillus sinesaloumensis | Litchfieldia sinesaloumensis |  |
| Candidatus Photodesmus katoptron | andidatus Photodesmus anomalopis |  |
| Lysinibacillus yapensis | Ureibacillus yapensis |  |
| Pseudalkalibacillus hwajinpoensis | Guptibacillus hwajinpoensis |  |
| Candidatus Nanopelagicus limnes | Candidatus Nanopelagicus limnae |  |
| Candidatus Enterovibrio escacola | Candidatus Enterovibrio escicola |  |
| Pseudomonas carboxydohydrogena | Afipia carboxydohydrogena |  |
| Kocuria indica | Kocuria marina |  |
| Candidatus Kuenenia stuttgartiensis | Candidatus Kuenenia stuttgartensis |  |
| Rhizobium smilacinae | Aliirhizobium smilacinae |  |
| Fournierella | Allofournierella |  |
| Candidatus Jidaibacter acanthamoeba | Candidatus Jidaibacter acanthamoebae |  |
| Pseudomonas marianensis | Stutzerimonas marianensis |  |
| Youhaiella tibetensis | Paradevosia tibetensis |  |
| Arenibacterium arenosum | Sedimentitalea arenosa |  |
| Trujillella endophytica | Trujillonella endophytica |  |
| Glaciibacter flavus | Orlajensenia flava |  |
| Microbacterium agarici | Paramicrobacterium agarici |  |
| Peptoniphilus coxii | Aedoeadaptatus coxii |  |
| Ruminococcus bicirculans |  |  |
| Candidatus Pelagibacter ubique | Candidatus Pelagibacter communis |  |
| Parageobacillus caldoxylosilyticus | Saccharococcus caldoxylosilyticus |  |
| Rhodococcus fascians | Rhodococcoides fascians |  |
| Prevotella oris | Segatella oris |  |
| Prevotella oralis | Hoylesella oralis |  |
| Prevotella buccae | Segatella buccae |  |
| Geobacillus thermoglucosidasius | Parageobacillus thermoglucosidasius |  |
| Bacteroides vulgatus | Phocaeicola vulgatus |  |
| Lactobacillus sakei | Latilactobacillus sakei |  |
| Lysobacter caseinilyticus | Noviluteimonas caseinilytica |  |
| Monaibacterium marinum | Pontivivens marinum |  |

**Supplementary Table 7. Phyla Uniquely detected by 16S rRNA and rpoB sequencing**

| **Target** | **Phylum** |
| --- | --- |
| *16S rRNA* | **Arcahea:** Nitrososphaerota, Methanobacteriota, and Euryarchaeota (sediment)  **Candidate Phyla:** Candidatus Dormiibacterota, candidate_division_NC10 and Candidatus Melainabacteria |
| *rpoB* | Eisenbacteria, Krumholzibacteriota (soil and sediment), and Methylomirabilota, Zixibacteria, Elusimicrobiota, Omnitrophota, and Lernaellota (sediment) |

**Supplementary Table 8. Correlation coefficients between DOC removal rates and transcripts abundances.**

| **Target** | **Time points** | **BAC** | **Correlation transcript abundance** | **Correlation transcription ratio** |
| --- | --- | --- | --- | --- |
| *16S rRNA* | 0h-48h | BAC 10 | -0.381 | 0.076 |
|  | 8h-72h |  | 0.649 | 0.924 |
| *rpoB* | 0h-48h |  | 0.909 | 0.843 |
|  | 8h-72h |  | 0.992 | 0.966 |
| *16S rRNA* | 0h-48h | BAC 20 | 0.472 | -0.250 |
|  | 8h-72h |  | 0.981 | 0.870 |
| *rpoB* | 0h-48h |  | 0.995 | 0.980 |
|  | 8h-72h |  | 0.982 | 0.980 |
